# *TET2* lesions enhance the aggressiveness of *CEBPA-*mutant AML by rebalancing *GATA2* expression

**DOI:** 10.1101/2023.03.28.534511

**Authors:** Elizabeth Heyes, Anna S. Wilhelmson, Anne Wenzel, Gabriele Manhart, Thomas Eder, Mikkel B. Schuster, Edwin Rzepa, Sachin Pundhir, Teresa D’Altri, Coline Gentil, Manja Meggendorfer, Jürg Schwaller, Torsten Haferlach, Florian Grebien, Bo T. Porse

## Abstract

The myeloid transcription factor CEBPA is recurrently biallelically mutated (i.e., double mutated; *CEBPA*^DM^) in acute myeloid leukemia (AML) with a combination of hypermorphic N-terminal mutations (*CEBPA*^NT^), promoting expression of the leukemia-associated p30 isoform, and amorphic C-terminal mutations. The most frequently co-mutated genes in *CEBPA*^DM^ AML are *GATA2* and *TET2*, however the molecular mechanisms underlying this co-mutational spectrum are incomplete.

By combining transcriptomic and epigenomic analyses of *CEBPA*-*TET2* co-mutated patients with models thereof, we identify *GATA2* as a conserved target of the *CEBPA*-*TET2* mutational axis, providing a rationale for the mutational spectra in *CEBPA*^DM^ AML. Elevated CEBPA levels, driven by *CEBPA*^NT^, mediate recruitment of TET2 to the *Gata2* distal hematopoietic enhancer thereby increasing *Gata2* expression. Concurrent loss of TET2 in *CEBPA*^DM^ AML induces a competitive advantage by increasing *Gata2* promoter methylation, thereby rebalancing GATA2 levels. Of clinical relevance, demethylating treatment of *Cebpa-Tet2* co-mutated AML restores *Gata2* levels and prolongs disease latency.

## INTRODUCTION

Acute myeloid leukemia (AML) is characterized by genetic alterations affecting proliferation and/or differentiation of hematopoietic stem or progenitor cells (HSPCs). Thereby, expansion of immature myeloid precursors, at the expense of normal hematopoiesis, ultimately leads to bone marrow (BM) failure if left untreated. Recent sequencing efforts have identified numerous recurrent mutations in AML and revealed patterns of mutational co-segregation, suggesting that synergism between certain lesions drives leukemogenesis^1^. While we now recognize these patterns, the mechanistic basis for context-specific positive or negative selection of certain lesions remains to be elucidated in most cases.

CCAAT enhancer binding protein alpha (CEBPA) is a hematopoietic lineage-specific transcription factor that binds and primes genes for myeloid development, and is required for differentiation and maturation of granulocytes^2^. The gene encoding CEBPA is biallelically mutated (i.e., double mutated; *CEBPA*^DM^) in 3–15% of *de novo* AML patients^3–9^. *CEBPA*^DM^ patients harbor either biallelic N-terminal mutations or a combination of a monoallelic N-terminal mutation together with a C-terminal mutation in the other allele. Whereas N-terminal CEBPA (*CEBPA*^NT^) lesions promote the expression of the truncated p30 isoform, C-terminal mutations result in CEBPA variants that are unable to dimerize or bind DNA, thus rendering them inactive. Hence, CEBPA p30 homodimers are the sole entity with functional transcription factor activity in *CEBPA*^DM^ AML. This is in contrast to normal hematopoietic cells where the full-length p42 isoform is predominantly expressed^2^. CEBPA p30 lacks two of three transactivation elements present in p42, but retains one transcriptional activating element and the basic-region leucine-zipper, which enables dimerization and DNA-binding^10^. CEBPA p30 has functions distinct from CEBPA p42 and can bind an isoform-specific set of enhancers and regulate the expression of downstream effector genes, such as *Nt5e* and *Msi2*^11, 12^. Importantly, in the context of *CEBPA*^DM^ AML, the *CEBPA*^NT^ is hypermorphic, leading to higher levels of the transcription factor, and thus, increased binding to enhancers and subsequent deregulation of gene expression^11^. In line with these data, mice with CEBPA p30 expression driven from the endogenous *Cebpa* locus develop AML with full penetrance within a year^13^.

Most patients with *CEBPA*^DM^ AML also feature additional mutations in *GATA2*, *TET2*, *WT1*, *NRAS*, *FLT3,* or *CSF3R*^9^. Several of these mutations are found together with *CEBPA^DM^* more frequently than expected by the individual frequency of each mutation, while other combinations are statistically underrepresented. Recent studies have shed light on the molecular mechanisms underlying mutational cooperativity for some of the co-mutated genes, *i.e. GATA2*^14^ and *CSF3R*^15^, while mechanistic insight is still lacking for other subgroups of *CEBPA*^DM^ AML. Of particular importance are mutations in the gene encoding the methylcytosine dioxygenase TET2 which, by converting 5-methylcytosine to 5-hydroxymethylcytosine, promotes DNA demethylation. *TET2* mutations (*TET2*^MUT^) are frequent in *CEBPA*^DM^ AML cases and are associated with inferior prognosis^16, 17^. Moreover, loss of *Tet2* has been implicated in accelerating and/or aggravating hematological malignancies in combination with several other recurrent gain-of-function and loss-of-function mutations^18–20^, reflecting the importance of appropriately regulated DNA demethylation in normal hematopoiesis. Importantly, while *Tet* loss alone only mildly affects hematopoiesis with myeloid skewing and increased competitiveness of HSCs^18^, as well as increased propensity of leukemic blasts to switch to a more stem-like phenotype^21^, it does not induce overt leukemia *per se*^22–24^. Despite being extensively studied, mechanistic insights of how TET2 loss-of-function cooperates with other aberrations has been hampered by the fact that malignant cells have been compared to their normal, wild-type counterparts in many studies.

In the present work, we sought to overcome this limitation by comparing *CEBPA-*mutant AML in the presence and absence of additional mutations in *TET2*. By combining transcriptomic and epigenomic analyses of relevant *in vitro* and *in vivo* models as well as data from AML patients, we identified an intricate mechanism where TET2 loss-of-function rebalances *Gata2* expression levels in *Cebpa*^DM^ AML, and hence drives an aggressive disease.

## RESULTS

### *TET2* mutations impair outcome for patients with *CEBPA*-mutant AML

To validate previous reports on the spectrum of co-occurring mutations in *CEBPA*^DM^ AML patients, we compiled data from 557 *CEBPA*^DM^ cases and evaluated the co-occurrence of other known leukemia driver mutations^3–7, 17^. *TET2* was the second most frequently co-mutated gene, with 1 in 5 *CEBPA*^DM^ cases harboring *TET2* mutations (**Figure 1A**; **Supplemental table 1**). Importantly, the survival of *TET2*-mutant (*TET2*^MUT^) *CEBPA*^DM^ patients was significantly lower than *TET2* wild-type (*TET2*^WT^) *CEBPA*^DM^ patients (**Figure 1B**), consistent with previous reports^16^, while the presence of *TET2* mutations did not cause a higher overall number of mutations in *CEBPA*^DM^ patients (**Supplemental figure 1A**). To investigate the functional consequences of *TET2* and *CEBPA* co-mutations, we analyzed RNA sequencing (RNA-seq) data from the Beat AML dataset^1^. We identified 1546 up- and 1201 downregulated genes in patients harboring a combination of *CEBPA* and *TET2* mutations when compared to *CEBPA*-mutant patients with wild-type *TET2* (**Figure 1C**). In line with the lower overall survival of *TET2*^MUT^*CEBPA*^DM^ patients, pathways related to inflammation, hypoxia, and aggressive cancer were upregulated in *CEBPA-TET2* co-mutated patients (**Supplemental figure 1B**). These findings indicate that mutations in *TET2* enhance the aggressiveness of *CEBPA*-mutant AML by deregulation of critical cellular pathways.

**Figure 1:**
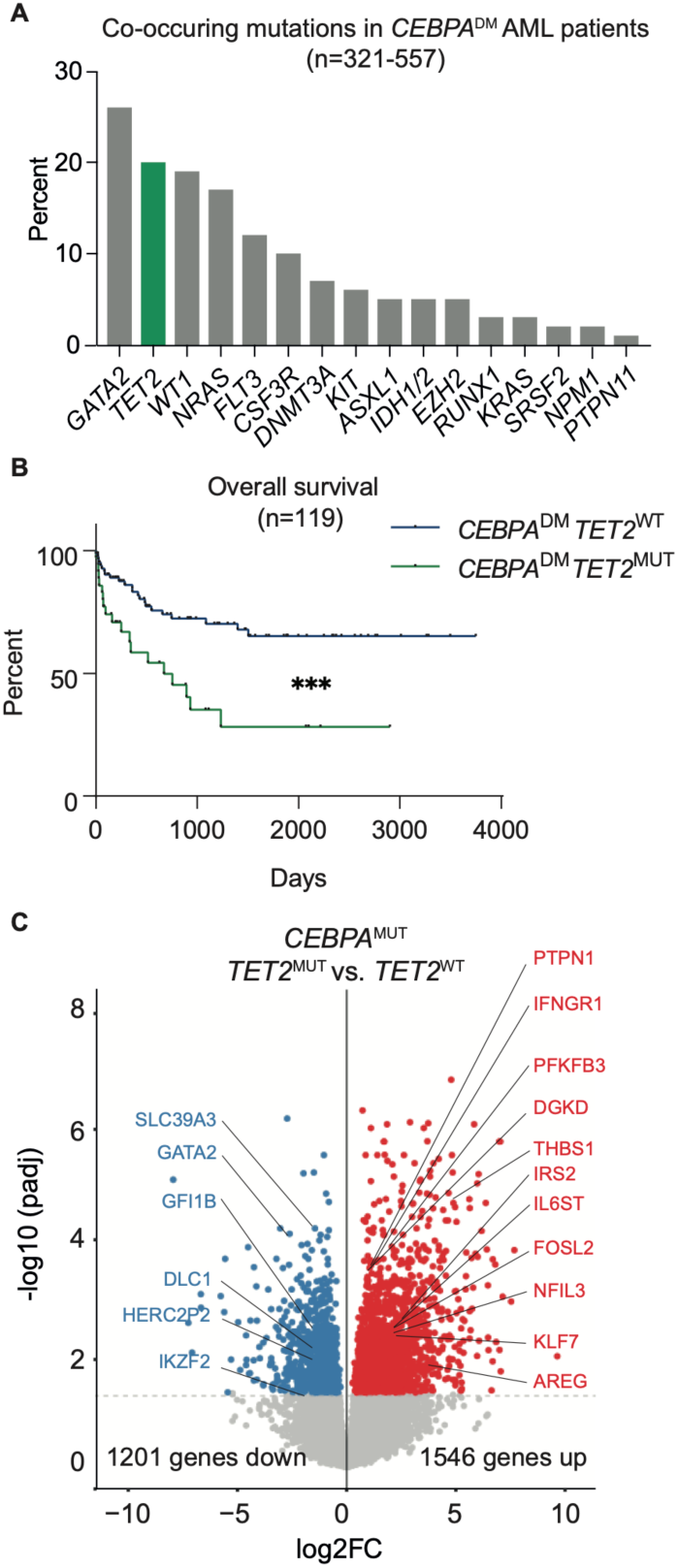
*TET2* mutations impair outcome for patients with *CEBPA*-mutant AML. (**A**) Frequency of co-occurring mutations in *CEBPA*^DM^ AML cases data aggregated from published cohorts^3–8, 17^ (n=321– 557). (**B**) Overall survival of *CEBPA*^DM^ patients with wild-type (*TET2*^WT^) or mutated TET2 (*TET2*^MUT^) (n=116). (**C**)Volcano plot depicting differentially expressed genes dependent on *TET2* mutational status in the cohort of *CEBPA*-mutant patients in the Beat AML dataset (n=5–11 per group). ***=P<0.001

### TET2 deficiency accelerates *Cebpa*-mutant AML

To study the effect of *TET2* mutations in *CEBPA*^DM^ AML in pathophysiologically relevant *in vitro* and *in vivo* models, we utilized cell and murine models in which expression of the p30 isoform is retained (*Cebpa*^p30/p30^ or *Cebpa*^−/p30^), while the normal p42 isoform of CEBPA is completely lost^13^. Since *TET2* is predominantly inactivated by loss-of-function mutations^25^, we modeled *TET2* mutations either by introduction of mutations with the CRISPR-Cas9 technology or by conditional knockout of the *Tet2* alleles.

First, we introduced *Tet2* mutations into a murine myeloid progenitor cell model (*Cebpa*^p30/p30^) (**Figure 2A**). *Tet2*-targeted cells displayed a selective advantage, as they outcompeted *Cebpa*^p30/p30^ cells (**Figure 2B**). Detailed analysis of the *Tet2* mutation that was associated with the proliferative advantage showed that the *Tet2* locus had acquired a +1 insertion in exon 3, which resulted in a downstream premature termination codon (**Supplemental figure 2A–B**). In line with this, clones isolated from the targeted cell pool exhibited strongly reduced TET2 protein expression (**Supplemental figure 2C**). Gene expression analysis revealed that *Tet2* loss in *Cebpa*^p30/p30^ cells caused downregulated expression of 916 genes, while only 474 genes were upregulated (**Figure 2C**). Gene set enrichment analysis (GSEA) showed higher expression of MYC and E2F targets in *Cebpa*^p30/p30^ *Tet2*-mutated cells, consistent with their proliferative advantage (**Supplemental figure 2D**).

**Figure 2:**
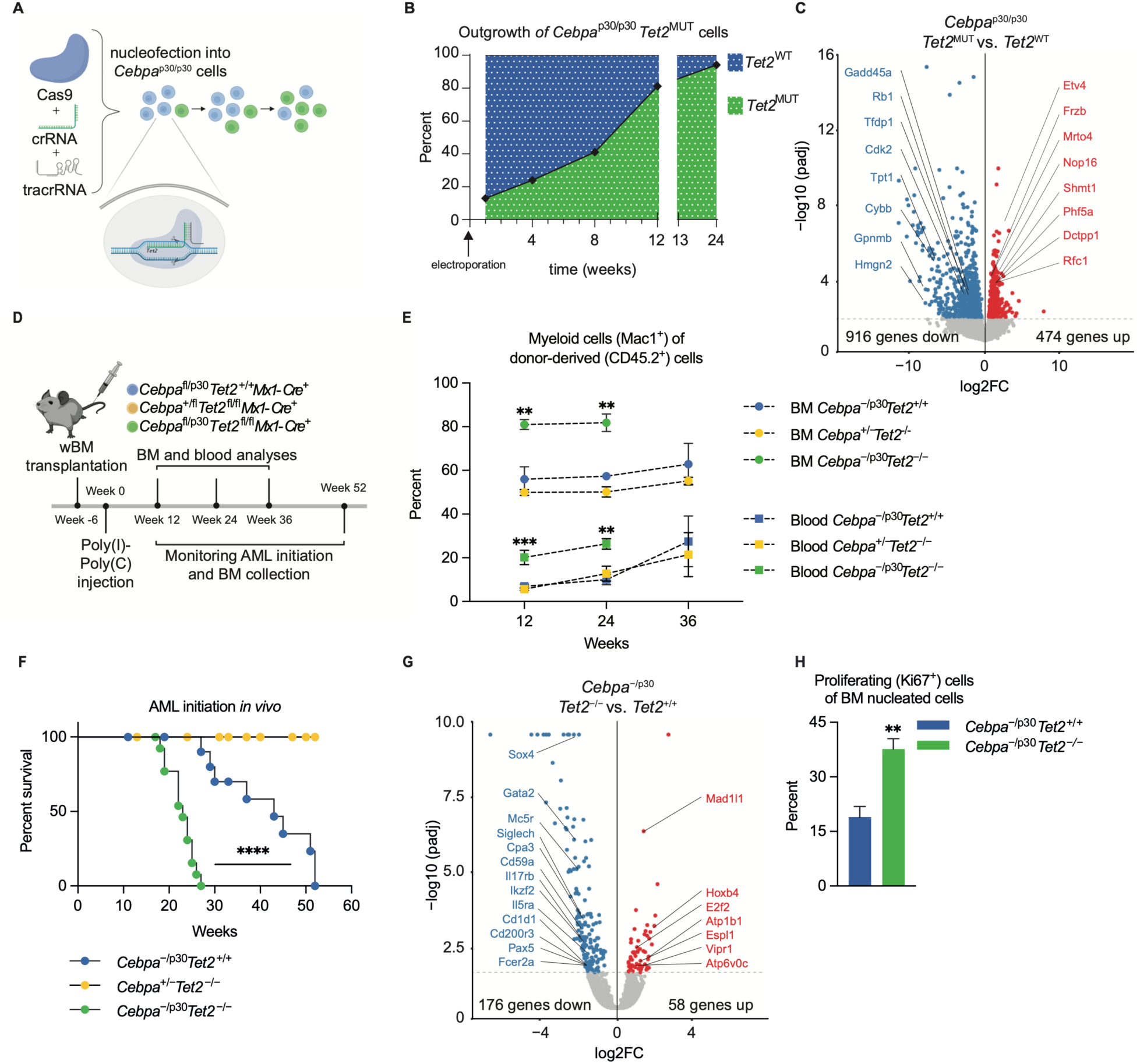
TET2 deficiency accelerates *Cebpa*-mutant AML. (**A**) Schematic representation of generation of *Tet2*-knockout clones with CRISPR/Cas9. (**B**) Proliferative outgrowth of *Cebpa*^p30/p30^ cells with *Tet2* indels. (**C**) Volcano plot depicting differentially expressed genes dependent on the *Tet2* mutational status in *Cebpa*^p30/p30^ cells (n=5–7 per group). (**D**) Experimental setup for evaluating the effect of *Tet2*-deficiency (*Tet2*^−/−^) in *Cebpa*^DM^ AML initiation *in vivo*. (**E**) Myeloid (Mac1^+^) contribution of donor-derived blood and bone marrow (BM) cells evaluated at 12, 24, and 36 weeks after BM transplantation and Cre-LoxP recombination to generate a *Cebpa*^−/p30^ and *Tet2*^−/−^ hematopoietic compartment (n=3–6 per genotype and timepoint). (**F**) Survival of lethally irradiated recipient mice after BM transplantation and Cre-LoxP recombination (n=12–14/group). (**G**) Volcano plot depicting differentially expressed genes dependent on *Tet2* deficiency status in *Cebpa*^−/p30^ leukemic blasts (n=3 per group). (**H**) Frequency of proliferating (Ki67^+^) cells in BM of moribund recipient mice (n=3 per group). **=P<0.01, ***=P<0.001, ****=P<0.0001

In summary, these data show that CRISPR/Cas9-induced TET2 loss provides a competitive advantage to myeloid progenitors expressing the oncogenic CEBPA variant p30.

Next, we wanted to assess the impact of hematopoietic expression of CEBPA p30 (*Cebpa*^−/p30^) with TET2-deficiency (*Tet2*^−/−^) on AML initiation *in vivo*. To do so, we transplanted lethally irradiated recipient mice with BM cells derived from mice with relevant allele combinations and, following hematopoietic reconstitution, induced hematopoietic-specific knockout of the *Cebpa* WT allele and/or the *Tet2* alleles (**Figure 2D**). The combination of CEBPA p30 expression with *Tet2* loss led to an early expansion of myeloid (Mac1^+^) cells in the BM and blood compared to mice with hematopoietic cells featuring either alteration on its own (**Figure 2E**). Conforming to patient data and data obtained from *Cebpa^p^*^30^*^/^*^30^ cells, *Cebpa*^−/p30^*Tet2*^−*/*−^ hematopoietic cells gave rise to AML with shorter latency than *Cebpa*^−/p30^*Tet2*^+/+^ cells, with a median survival of 23 and 43 weeks, respectively (**Figure 2F**). TET2 deficiency alone (*Cebpa*^+/−^*Tet2*^−/−^) did not give rise to AML and cells which retained expression of the p42 isoform from one allele (*Cebpa*^+/p30^) only sporadically underwent leukemic transformation (**Figure 2F**; **Supplemental figure 2E**). The transformed blasts expressed myeloid (Mac1^+^) and granulocytic (Gr1^+^) markers, confirming myeloid origin of the leukemia (**Supplemental figure 2F**). The leukemias were transplantable into secondary recipients, and the shorter latency of the TET2-deficient *Cebpa*^DM^ AML was preserved in this setting (**Supplemental figure 2G–H**), indicating that TET2 not only has important tumor suppressive functions during malignant transformation but also during progression of AML.

We performed RNA-seq on *Cebpa*^−/p30^ (*Tet2* WT and knockout) AML blasts to assess changes in gene expression upon TET2 deficiency. Again, we found that the majority of differentially expressed genes was decreased in TET2-deficient AML blasts, with 176 down- vs. 58 upregulated genes (**Figure 2G)**. GSEA highlighted upregulation of genes involved in IL-6-JAK-STAT-signaling and hypoxia, in line with RNA-seq data from human *TET2*^MUT^*CEBPA*^MUT^ cases (**Supplemental figure 1B**; **Supplemental figure 2I**). Furthermore, pathways related to cell cycle progression (G2M checkpoint and E2F targets) were enriched in TET2-deficient AML, indicating increased growth upon loss of TET2, consistent with the effects observed in the cell model (**Supplemental figure 2D**; **Supplemental figure 2I**). In line with this, we found that a higher frequency of *Cebpa*^−/p30^*Tet2*^−/−^ blasts expressed the proliferation marker Ki67 (**Figure 2H**). In addition, we also observed increased proliferative capacity of *Cebpa*^−/p30^*Tet2*^−/−^ blasts compared to *Cebpa*^−/p30^*Tet2*^+/+^ blasts *ex vivo*. This difference was dependent on *Tet2* status, as the TET2 co-factor Vitamin C was able to mitigate proliferation of *Cebpa*^−/p30^*Tet2*^+/+^ but not of *Cebpa*^−/p30^*Tet2*^−/−^ cells (**Supplemental figure 2J**).

Collectively, these data show that TET2 deficiency accelerates the establishment and progression of CEBPA p30-driven AML *in vivo*.

### Loss of TET2 leads to reduced *Gata2* levels in *Cebpa*-mutant AML

To find conserved gene targets of the CEBPA-TET2 axis, we integrated the transcriptomic data from our *in vitro* and *in vivo* models with gene expression analyses from AML patients harboring *CEBPA* and *TET2* mutations. Three target genes exhibited downregulated expression in all three data sets; *FUT8*, *GATA2*, and *SIRT5* (**Figure 3A**; **Supplemental figure 3A–C**).

**Figure 3:**
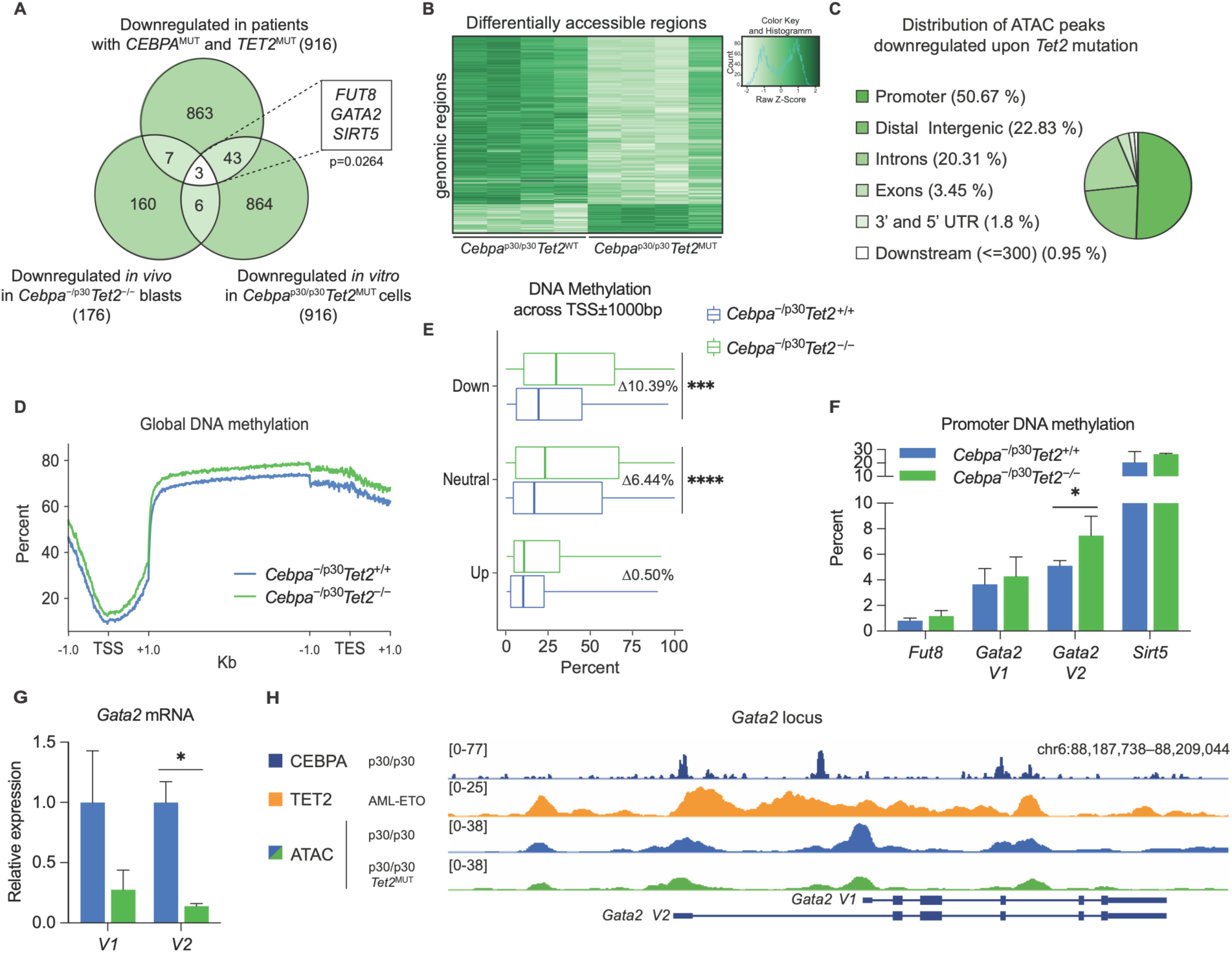
Loss of TET2 leads to reduced *Gata2* levels in *Cebpa*-mutant AML. (**A**) Conserved targets of the CEBPA-TET2 axis visualized in a Venn-diagram of downregulated genes in *CEBPA*-*TET2* co-mutated AML overlaid with corresponding data from *Tet2*-deficient *in vivo* and *in vitro* models of *Cebpa*^DM^ AML (P=0.0264 vs. number of overlapping genes expected by random distribution). (**B**) Heatmap of differentially accessible regions assessed by assay for transposase-accessible chromatin sequencing (ATAC-seq; |Log2FC|>1, p<0.01), and (**C**) genomic distribution of downregulated peaks (FDR < 0.05) upon *Tet2* mutation (n=4/group). (**D**) Representative genome wide DNA-methylation status assessed by whole genome bisulfite sequencing (WGBS) showing frequency of methyl-cytosine (mC) across the transcription start site (TSS) ±1000 base pairs, gene body scaled to 4000 base pairs, and transcription termination site (TES) ±1000 base pairs. (**E**) Median and interquartile range of percent mC at promoters of up-, non- and down-regulated genes (n=2–3/group). Whiskers indicates max–min. (**F**) Promoter DNA methylation of conserved target genes in *Cebpa*^−/p30^*Tet2*^−/−^ and *Cebpa*^−/p30^*Tet2*^+/+^ leukemic blasts (n=2–3/group). (**G**) *Gata2* variant mRNA expression in *Cebpa*^−/p30^*Tet2*^−/−^ and *Cebpa*^−/p30^*Tet2*^+/+^ leukemic blasts (n=3/group). (**H**) Schematic genomic view of the *Gata2* locus, including tracks from CEBPA chromatin immunoprecipitation sequencing (ChIP-seq) in *Cebpa*^p30/p30^ cells (data from Heyes et al.^12^), TET2 ChIP-seq in AML-ETO expressing cells (data from Rasmussen et al.^26^), and ATAC-seq in *Cebpa*^p30/p30^ cells. *=P<0.05, ***=P<0.001, ****=P<0.0001

Although the deregulation of these three genes was observed across species and differential experimental setups, we next aimed to investigate if their decreased gene expression was a direct result of TET2 deficiency. We therefore assessed chromatin accessibility and DNA methylation as a proxy for TET2 binding and activity ^26^. Through assay for transposase-accessible chromatin sequencing (ATAC-seq), we identified 2552 differentially accessible regions in *Tet2*^MUT^*Cebpa*^p30/p30^ vs. *Tet2*^WT^*Cebpa*^p30/p30^ cells, and consistent with an activating effect of TET2, the majority of differential regions were less accessible in TET2-deficient cells (**Figure 3B**). Half of the ATAC-seq peaks downregulated upon *Tet2* mutation were located in promoters, and these regions were enriched for GATA and NFAT motifs (**Figure 3C**; **Supplemental figure 3D**). Using whole genome bisulfite sequencing (WGBS), we observed a global increase in DNA methylation in *Cebpa*^−/p30^*Tet2*^−/−^ vs. *Cebpa*^−/p30^*Tet2*^+/+^ AML blasts, consistent with a loss of demethylase activity in *Tet2* knockout blasts (**Figure 3D**). Increased DNA methylation was observed in promoter regions of genes whose expression were downregulated upon TET2 loss, while upregulated genes did not show any changes (+54%; **Figure 3E**). Strikingly, this pattern was not apparent when DNA methylation was evaluated across gene bodies (**Supplemental figure 3E**). Non-regulated, neutral genes exhibited equal increase in DNA methylation across promoter and gene body (**Figure 3E**; **Supplemental figure 3E**). Thus, loss of TET2 in *Cebpa*^DM^ cells caused decreased chromatin accessibility and increased methylation of DNA in promoters of TET2-responsive genes, consistent with previous reports showing that TET2 binding is enriched in promoters of TET2-regulated genes^27^.

To identify direct CEBPA-TET2 gene target(s), we evaluated the previously identified conserved candidates based on changes in DNA methylation of their promoters. Out of the three target genes, only the gene encoding the transcription factor GATA-binding factor 2 (GATA2) showed a gain of DNA methylation in the promoter of the gene variant 2 (*Gata2 V2*) upon TET2 deficiency (+46%; **Figure 3F**). In line with this, specifically the Gata*2 V2* mRNA isoform was downregulated in TET2-deficient *Cebpa*^DM^ AML blasts (−86%; **Figure 3G**), while changes in mRNA expression and promoter methylation of *Gata2 V1* did not reach statistical significance (**Figure 3F–G**).

In summary, these analyses identify *Gata2* (locus overview in **Figure 3H**) as a conserved target of the CEBPA-TET2 axis across several settings. TET2 deficiency causes increased DNA methylation of the *Gata2* promoter, resulting in reduced mRNA expression.

### Moderate *Gata2* reduction increases competitiveness of *Cebpa-*mutant AML

GATA2 is an essential transcription factor for hematopoietic cells and has profound effects on HSC maintenance. Moreover, it is recurrently mutated in AML^28, 29^ and *GATA2* lesions are overrepresented in *CEBPA*^DM^ AML^8, 16, 30–32^. Given these critical roles of GATA2, we next examined the consequences of reduced GATA2 levels in *CEBPA*^DM^ AML. To test if reduced *Gata2* expression would provide a competitive advantage *in vivo*, we set up an RNA-interference (RNAi) based competition assay (**Figure 4A**) utilizing established *Cebpa*^p30/p30^ leukemia cells. First, we identified four short hairpin RNAs (shRNA) which lowered *Gata2* expression to a varying degree (**Figure 4B**). Upon transplantation of shRNA-expressing cells we observed a non-monotonic relationship between *Gata2* expression levels and competitiveness, as measured by sh*Gata2*-to-shControl ratios. While efficient downregulation of *Gata2* expression (>75%) did not provide any competitive advantage to *Cebpa*^DM^ cells, moderate silencing (25–60%) imposed a three-fold increase in their ability to compete (**Figure 4C–D**). Repetition of this experiment including only the most and least efficient shRNAs in a separate experiment yielded similar results (**Supplemental figure 4A–B**). To test if the same effects are observed in an *in vitro* setting, we targeted *Gata2* in *Cebpa*^p30/p30^ cells using the CRISPR/Cas9 approach. *Gata2*-targeted cells showed a proliferative advantage over *Gata2*^WT^ cells, leading to their outgrowth (**Figure 4E–F**). In accordance with previously published data that complete loss of *Gata2* expression results in a loss of competitiveness^33–,35^, we found that only clones with heterozygous *Gata2* inactivation were viable, while clones with homozygous mutations in *Gata2* could not be recovered (**Supplemental figure 4C**).

**Figure 4:**
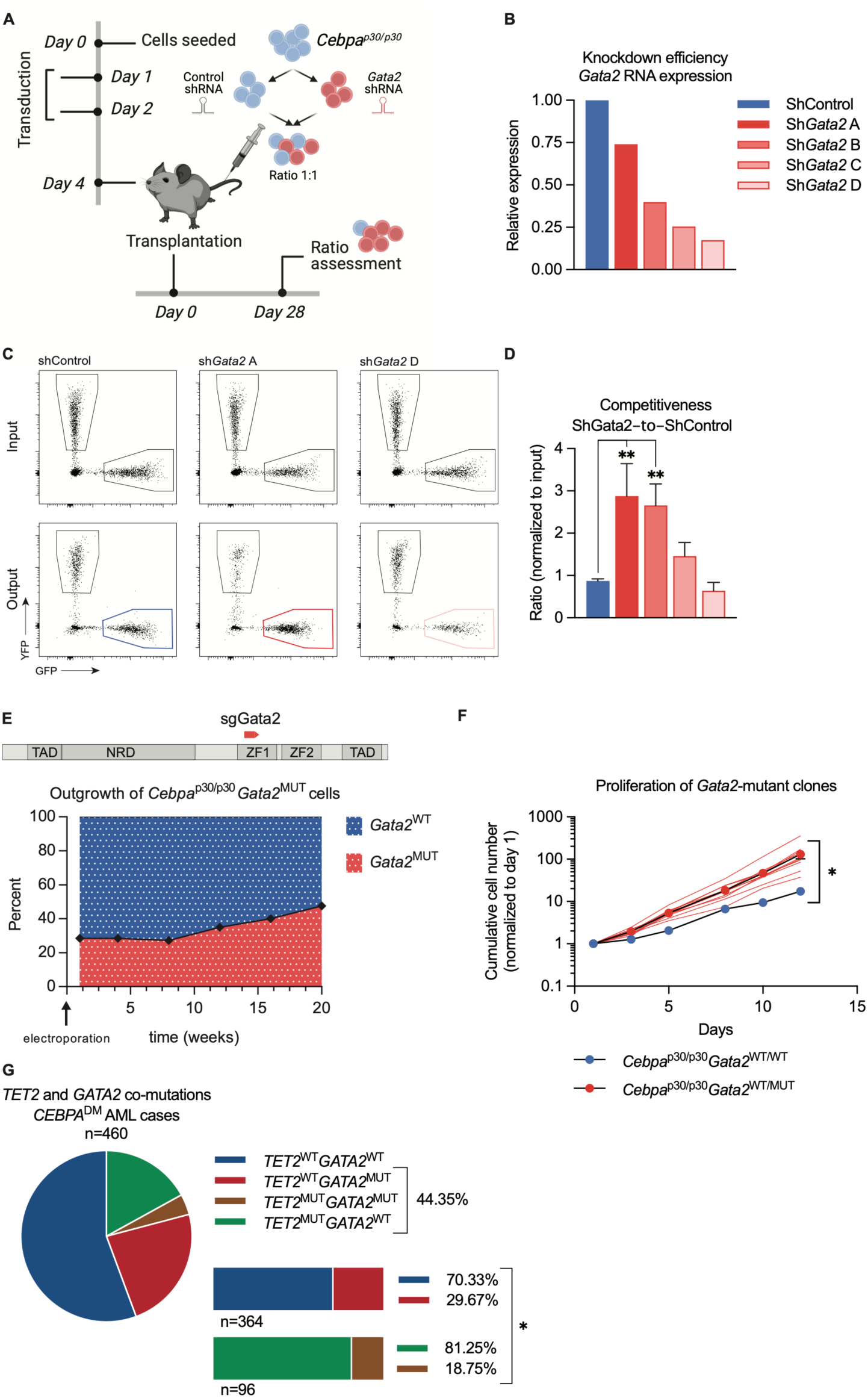
Moderate *Gata2* reduction increases competitiveness of *Cebpa-*mutant AML. (**A**) Experimental setup for evaluating the effect of *Gata2* knockdown, via short hairpin RNA (shRNA) mediated silencing, on *Cebpa*^p30/p30^ leukemic cells in a competitive *in vivo* assay. (**B**) *Gata2* mRNA in *Cebpa*^p30/p30^ leukemic cells prior to transplantation. (**C**) Representative flow cytometry profiles of input and output of shControl (no knockdown), sh*Gata2*A (low knockdown), and sh*Gata2*D (high knockdown). (**D**) Competitive advantage of targeting shRNA (GFP^+^) vs. non-targeting shRNA (YFP^+^) cells *in vivo* assessed as by flow cytometry (n=3–4 per group). (**E**) Experimental setup for *Gata2* CRISPR/Cas9 mutagenesis in *Cebpa*^p30/p30^ cells, and outgrowth of heterozygous mutated clones. Percentages of *Gata2* mutated clones are indicated. (**F**) Growth curve of *Cebpa*^p30/p30^ clones with *Gata2* mutation (*Cebpa*^p30/p30^*Gata2*^+/MUT^) or wild type *Gata2* (*Cebpa*^p30/p30^*Gata2*^+/+^). Red lines mark individual clones. **(G)** Presence or absence of *GATA2* mutations (*GATA2*^MUT^) in *CEBPA* double mutated (*CEBPA*^DM^) AML cases (n=460) with or without *TET2* mutations (*TET2*^MUT^) in aggregated data from published cohorts^3–5, 7, 8, 17^. *=P<0.05, **=P<0.01

If the pro-leukemogenic effect of *TET2* mutations was, at least partly, caused by lowering *GATA2* expression, we reasoned that concomitant mutations in both genes would be redundant and thus, the pattern of TET2 and GATA2 mutations would be mutually exclusive. Indeed, *TET2*^MUT^*CEBPA*^DM^ AML cases showed a lower frequency of *GATA2* mutations than expected from the frequency of *GATA2* mutations in *TET2*^WT^*CEBPA*^DM^ AML cases (**Figure 4G**; **Supplemental table 2A**), which was also true for all AML cases (3.5% in TET2^MUT^ vs. 8.9% in TET2^WT^; **Supplemental figure 4D**; **Supplemental table 2B**). Importantly, whereas mutations in *WT1* followed the same pattern as *TET2*, *CSF3R* mutations appeared in equal frequency between *TET2*^MUT^*CEBPA*^DM^ and *TET2*^WT^*CEBPA*^DM^ AML cases, and *ASXL1* mutations were increased in *TET2*^MUT^*CEBPA*^DM^ AML (**Supplemental figure 4E**; **Supplemental tables 2C-E**).

Altogether, our data suggest that loss of TET2 in *Cebpa*^DM^ AML causes a moderate decrease in *Gata2* expression, which in turn increases the competitive fitness of the leukemia. Hence, this indicates that *TET2* and *GATA2* mutations are partially redundant in *CEBPA*^DM^ AML, providing a mechanistic rationale for the mutational spectra observed in this AML entity.

### Increased CEBPA p30 binding to the *Gata2* distal hematopoietic enhancer drives expression of *Gata2* via TET2

We next asked if *GATA2* expression is dependent on *CEBPA* mutational status. To this end, we exploited published transcriptomics data from human and mouse *CEBPA*^DM^ AML^11^. *GATA2* expression was increased in human *CEBPA*^DM^ leukemic granulocyte/monocyte progenitors (GMPs) compared to GMPs from healthy donors (+77%; **Supplemental figure 5A**). Correspondingly, *Gata2* was upregulated in murine *Cebpa^p^*^30^*^/p^*^30^ leukemic GMPs as compared to normal GMPs (+43%; **Figure 5A**). Since CEBPA is known to exert its transcription factor activity by binding to enhancers and thereby promote gene expression^36^, we assessed binding of CEBPA to the crucial *Gata2* distal hematopoietic enhancer (*G2*DHE; −77 kb in mouse) that governs *Gata2* expression in hematopoietic stem and progenitor cells including GMPs^11, 37^. Notably, we found substantially increased levels of CEBPA bound to the *G2*DHE in *Cebpa*^p30/p30^ leukemic GMPs compared to normal counterparts (+147%; **Figure 5B**), while the binding levels associated with other known proximal and distal *cis-*regulatory elements of the *Gata2* gene were unchanged (**Supplemental figure 5B–C**). However, DNA methylation in the *G2*DHE was low and unaltered upon *Tet2* loss (**Supplemental figure 5D**). Importantly, CEBPA binding, as assessed by ChIP-qPCR, was unchanged upon introduction of *Tet2* mutations in *Cebpa*^p30/p30^ cells (**Supplemental figure 5E**).

**Figure 5:**
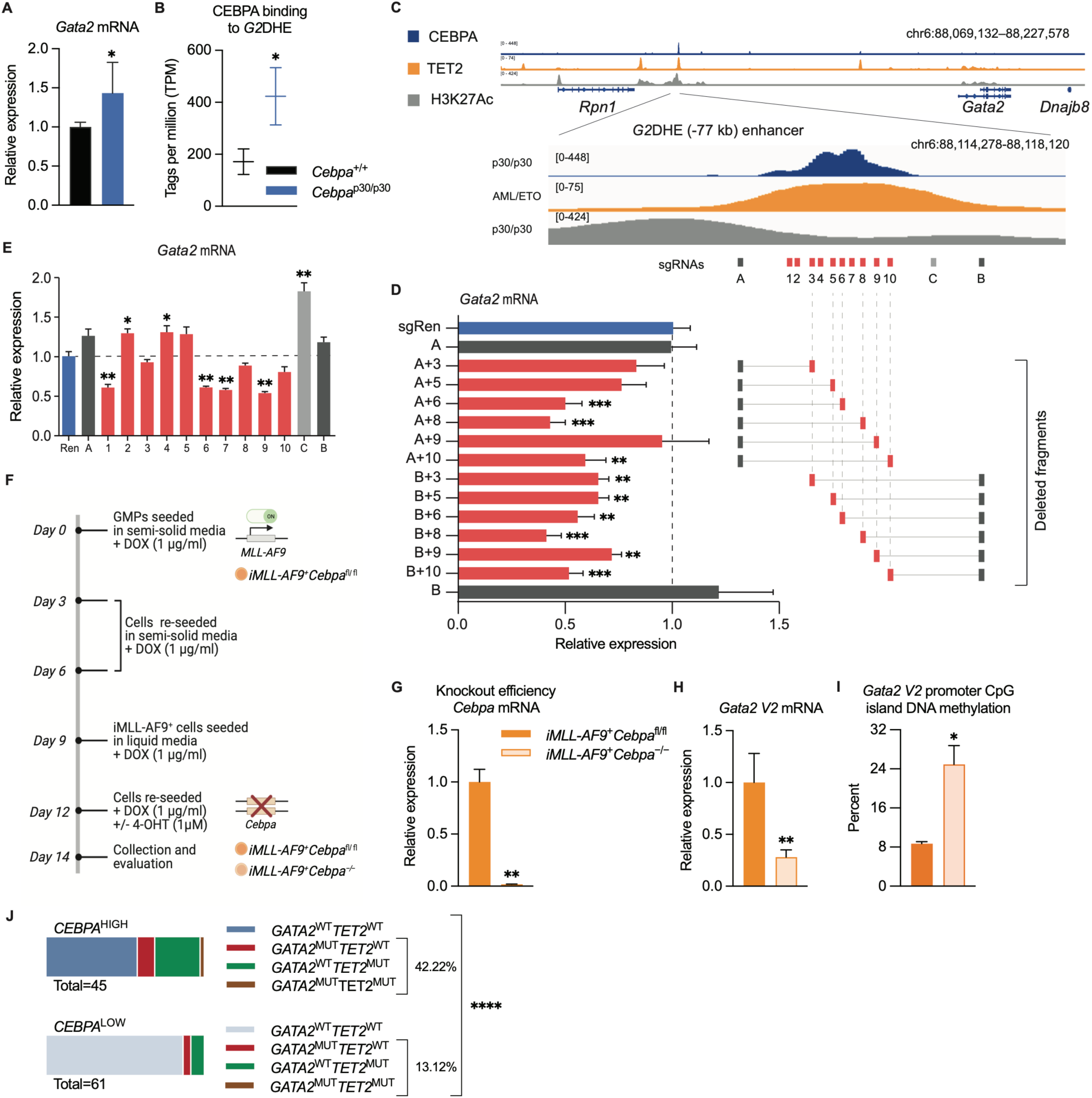
Increased CEBPA p30 binding to the *Gata2* distal hematopoietic enhancer drives expression of *Gata2* via TET2. (**A**) *Gata2* mRNA expression in mouse *Cebpa*^p30/p30^ leukemic granulocyte/monocyte progenitors (GMPs) vs normal GMPs and, (**B**) CEBPA binding to the *Gata2* distal hematopoietic enhancer (*G2*DHE; −77kb) region, data from Jakobsen et al.^11^ (n=2–4 per group). (**C**) Schematic genomic view of the *Gata2* distal hemeatopoietic enhancer (*G2*DHE), including tracks from CEBPA and H3K27Ac chromatin immunoprecipitation sequencing (ChIP-seq) in *Cebpa*^p^^30/p30^ cells (data from Heyes et al.^12^), TET2 ChIP-seq in AML-ETO expressing cells (data from Rasmussen et al.^26^),Targeting of the *G2*DHE by dual-(**D**) or single-(**E**) guided CRISPR-Cas9 in *Cebpa*^p30/p30^ cells using indicated sgRNAs (n=3/condition). (**F**) Experimental setup for evaluating the effects of *Cebpa* knockout on *Gata2 V2* mRNA expression and DNA methylation of the CpG island at the promoter of *Gata2 V2* in MLL-fusion driven AML (*iMLL-AF9*). (**G**) *Cebpa* and (**H**) *Gata2 V2*mRNA expression upon induction of Cre-LoxP recombination and, (**I**) DNA methylation of the *Gata2* V2 promoter CpG-island (2 biological replicates per genotype). (**J**) Frequency of *GATA2* and/or *TET2* mutations (*GATA2*^MUT^ and TET2^MUT^, respectively) in *CEBPA* high expressing (*CEBPA*^HIGH^ n=45) vs. *CEBPA* low expressing (CEBPA^LOW^ n=61) AML cases, data from Beat AML cohort^1^. *=P<0.05, **=P<0.01, ***=P<0.001, ****=P<0.0001

These results prompted us to test if CEBPA binding to the *G2*DHE modulates *Gata2* expression in *Cebpa*^DM^ AML. First, we deleted 250–500 bp fragments of the *Gata2* enhancer encompassing the CEBPA binding site using the CRISPR/Cas9 approach. *Gata2* mRNA expression was decreased upon targeting the genomic region with strong CEBPA binding compared to non-targeting sgRNAs or areas flanking the CEBPA-bound region (**Figure 5C–D**). Second, CRISPR/Cas9-mediated introduction of indels within 200 bp of the CEBPA consensus motif using several sgRNAs reduced *Gata2* expression (**Figure 5E**). Combined, these data suggest that CEBPA binding to the *G2*DHE is important for promoting *Gata2* expression in *Cebpa*^DM^ AML. Further, the *G2*DHE has been shown to primarily regulate expression of the hematopoietic specific *Gata2 variant 2* (*V2*)^38, 39^, conforming with our data that particularly the *Gata2 V2* promoter displayed an increase in DNA methylation and that the Gata*2 V2* mRNA was downregulated in TET2-deficient *Cebpa*^DM^ AML blasts (**Figure 3F-G**).

Next, we tested if reduction of CEBPA in AML cells influenced expression and promoter DNA methylation of *Gata2 V2*. Given the dependence of *CEBPA*^DM^ AML on CEBPA for survival and maintenance, we utilized MLL-fusion driven AML, in which CEBPA is dispensable for the maintenance of established leukemia^40^. Cre-mediated loss of *Cebpa* in leukemic cells expressing the inducible MLL-AF9 fusion-protein (*iMLL-AF9*^+^*Cebpa*^−/−^; **Figure 5F–G**) caused reduced *Gata2 V2* mRNA levels compared to control cells (*iMLL-AF9*^+^*Cebpa*^fl/fl^) (−72%; **Figure 5H**). Importantly, the methylation frequency of the CpG island located at the *Gata2 V2* promoter was increased in two separate leukemic lines (+186%; **Figure 5I**; **Supplemental figure 5F**), suggesting that *Gata2 V2* mRNA expression is regulated via the CEBPA-TET2 axis.

In light of these findings, we asked whether elevated CEBPA level, and not the *CEBPA* mutation(s) *per se*, drives the selective pressure for *GATA2-* and/or *TET2* loss in AML to achieve moderate *GATA2* levels that are optimal for leukemia growth. We therefore stratified AML cases in the Beat AML cohort^1^ based on *CEBPA* expression and assessed their *GATA2* and *TET2* mutational status. Indeed, the frequency of *GATA2* and/or *TET2* mutations was three-fold higher in *CEBPA*^HIGH^ AML compared to the *CEBPA*^LOW^ samples (**Figure 5J**). In line with previous data showing a hypermorphic effect of *CEBPA*^DM^ ^11^, the *CEBPA*^HIGH^ group contained the majority of the *CEBPA*-mutant cases in the cohort (82 and 100% of *CEBPA*^SM^ and *CEBPA*^DM^, respectively), while none of the cases in the *CEBPA*^LOW^ group were *CEBPA*-mutated. In conclusion, our data suggest that elevated CEBPA binding to the *G2*DHE, driven by the hypermorphic effect of *Cebpa*^NT^, increases TET2-mediated demethylation of the *Gata2* promoter, which leads to elevated *Gata2* levels in *Cebpa*^DM^ AML. In this context, *Cebpa*^DM^ AML cells gain a competitive advantage by loss of TET2, which in turn promotes an increase in DNA methylation at the *Gata2* promoter resulting in the rebalancing of *Gata2* levels.

### Demethylating treatment restores *Gata2* expression and prolongs survival in TET2-deficient *Cebpa*-mutant AML

Finally, we investigated if treatment with the demethylating agent 5-azacytidine (5-AZA) would be beneficial in TET2-deficient *CEBPA*^DM^ AML. *Ex vivo* treatment with 5-AZA restored *Gata2* expression in *Cebpa*^−/p30^*Tet2*^−/−^ blasts to levels observed in *Cebpa*^−/p30^*Tet2*^+/+^, while 5-AZA treatment did not affect *Gata2* levels in *Cebpa*^−/p30^*Tet2*^+/+^ cells (**Supplemental figure 6A**). Moreover, 5-AZA decreased the viability of blasts from both genotypes, although to a higher degree in the TET2-deficient setting (−82% and −40%, respectively, p<0.01; **Supplemental figure 6B**).

To evaluate if the enhanced effect of 5-AZA treatment in TET2-deficient AML would also hold true *in vivo*, mice were transplanted with *Cebpa*^−/p30^*Tet2*^−/−^ or *Cebpa*^−/p30^*Tet2*^+/+^ AML blasts and treated with 5-AZA for three consecutive days after disease establishment (**Figure 6A**). While the blast frequency of TET2-deficient *Cebpa*^−/p30^ AML decreased upon 5-AZA treatment (−87%; **Figure 6B**), the treatment did not significantly decrease the frequency of TET2-proficient cells. Furthermore, 5-AZA treatment restored *Gata2* levels in *Cebpa*^−/p30^*Tet2*^−/−^ blasts *in vivo* to the same level as in *Cebpa*^−/p30^*Tet2*^+/+^ blasts (**Figure 6C**). Importantly, a longer intermittent 5-AZA treatment prolonged the survival of mice transplanted with *Cebpa*^−/p30^*Tet2*^−/−^ blasts (median survival +22%; **Figure 6D-E**), while it did not affect disease latency of mice transplanted with *Cebpa*^−/p30^*Tet2*^+/+^ blasts.

**Figure 6:**
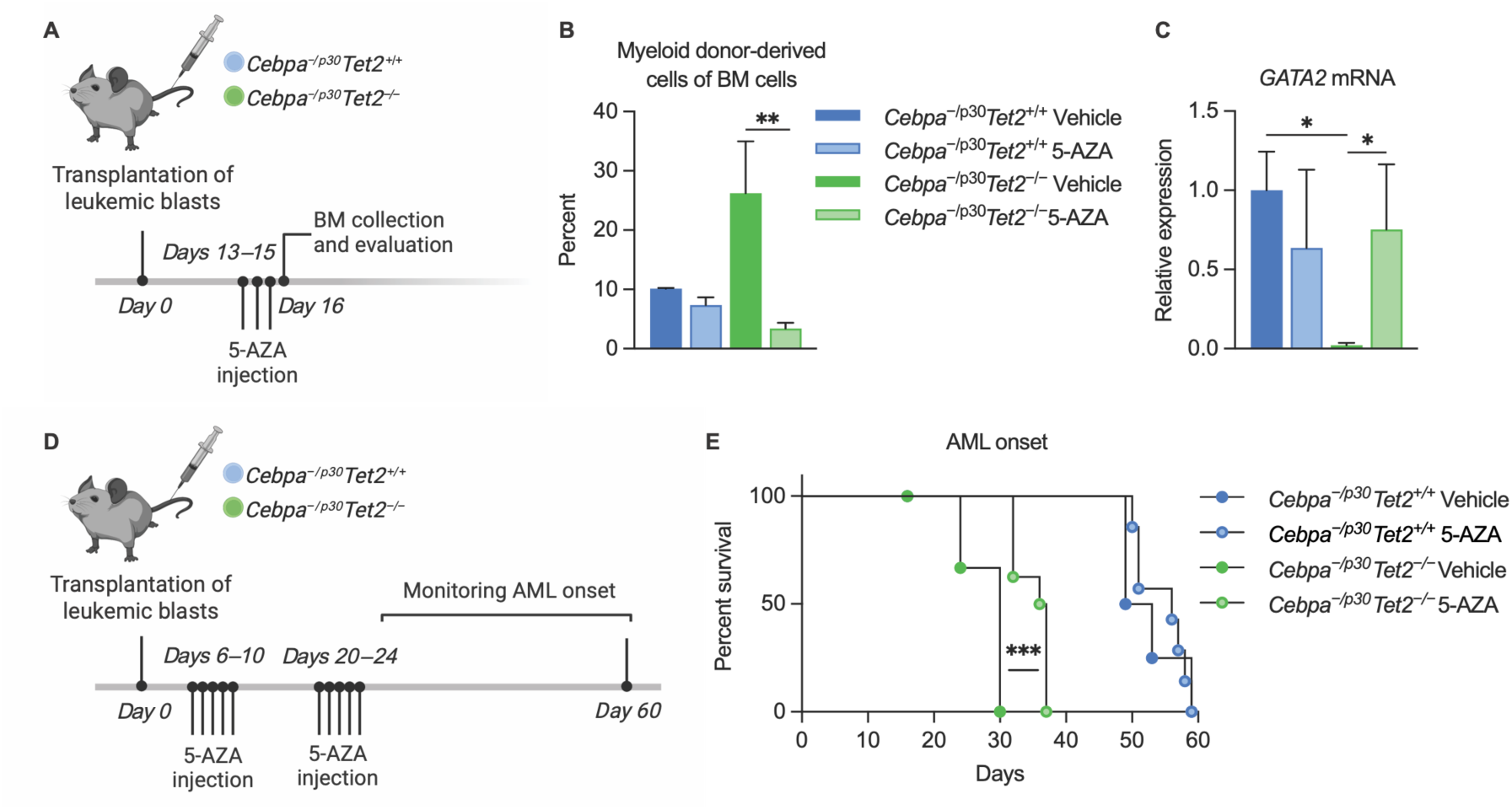
Demethylating treatment restores *Gata2* expression and prolongs survival in TET2-deficient *Cebpa*-mutant AML. (**A**) Experimental setup for evaluating the effect of short-term 5-azacytidine (5-AZA) treatment *in vivo*. (**B**) Expansion of myeloid (Mac1^+^) donor-derived cells in bone marrow (BM) assessed by flow cytometry, and (**C**) *Gata2* mRNA expression in sorted leukemic blasts by qPCR assessed 24 hours after the last of three injections of 5-AZA or vehicle (n=3–4 per group). (**D**) Experimental setup for evaluating the effect of 5-AZA treatment on AML progression *in vivo*. (**E**) Survival of sub-lethally irradiated tertiary recipient mice after transplantation of leukemic BM from moribund secondary recipient mice in response to intermittent 5-AZA treatment (5-AZA treated groups n=8 and vehicle treated groups n=4). *=P<0.05, **=P<0.01, ***=P<0.001

In summary, we show that the demethylating agent 5-AZA can restore *Gata2* expression levels in TET2-deficient *Cebpa*^DM^ AML to that of TET2-proficient *Cebpa*^DM^ AML, and concomitantly reduce leukemic burden and prolong survival of mice transplanted with TET2-deficient *Cebpa*^DM^ leukemic blasts.

## DISCUSSION

Mutational cooperativity is a fundamental driver of cancer development, progression, and aggressiveness. For *CEBPA*^DM^ AML, co-occurring lesions have been found in genes such as *GATA2*, *TET2*, *WT1*, *FLT3,* and *CSFR3*. While the mechanistic basis for the cooperation between *CEBPA* and *GATA2/CSFR3* mutations has been investigated using mouse models^14, 15^, we have very little insights into why other lesions, such as those in *TET2*, are overrepresented in *CEBPA*^DM^ AML. Here, we show that TET2 loss-of-function in *CEBPA*^DM^ AML leads to an aggressive disease phenotype by rebalancing the increased and suboptimal levels of *GATA2* that are induced by hypermorphic *CEBPA*^NT^ mutations driving CEBPA-p30 isoform expression (see model in **Figure 7A**). Specifically, loss of TET2 binding to the hematopoietic specific *G2*DHE enhancer results in increased DNA methylation in the promoter region of the hematopoietic-specific *Gata2* isoform (*Gata2 V2*). This proleukemic effect of TET2 loss can be reversed by the demethylating agent 5-azacytidine, suggesting that this could be a potential treatment option in *CEBPA*^DM^*TET2*^MUT^ patients. Altogether, our work proposes that CEBPA-mutant AMLs acquire additional lesions in genes such as *GATA2* and *TET2* to reestablish balanced *GATA2* levels that permit leukemia development and progression.

**Figure 7:**
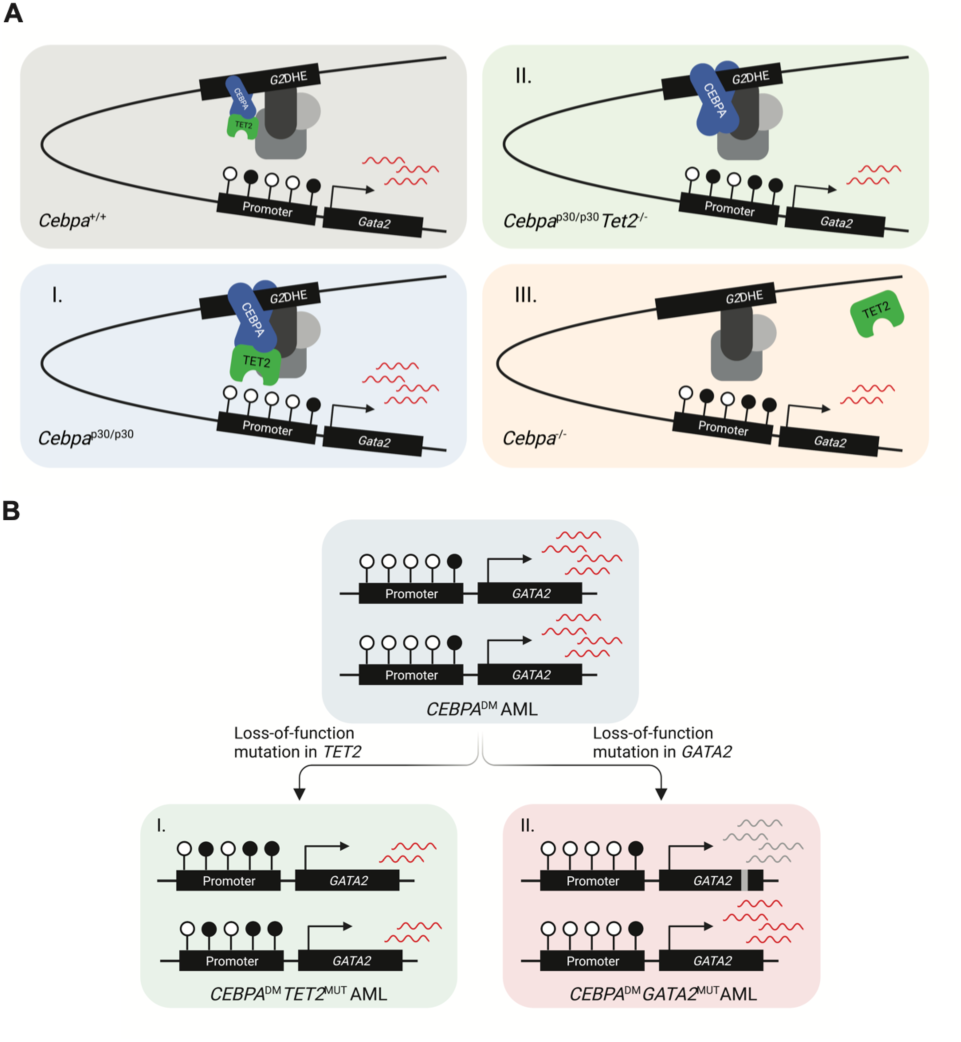
*TET2* lesions enhance the aggressiveness of *CEBPA-*mutant AML by rebalancing *GATA2* expression. (**A**) Model of *Gata2* differential expression as a consequence of (I) elevated CEBPA p30 due to the hypermorphic effect of the *CEBPA*^NT^, (II) TET2 deficiency and, (III) CEBPA deficiency. (**B**) Schematic illustration of two strategies for *CEBPA*^DM^ AML to rebalance *GATA2* levels by (I) loss-of-function mutations in *TET2* and (II) loss-of-function mutations in one *GATA2* allele.

Our work highlights the central importance of GATA2 regulation in *CEBPA*-mutant AML. Specifically, we show that *GATA2* is a conserved target gene of CEBPA and TET2. Furthermore, the elevated levels of the CEBPA p30 variant likely mediate *GATA2* upregulation in CEBPA-mutant AML. The increased expression of *Gata2* is counteracted by loss of *TET2* in *in vitro* and *in vivo* models of *Cebpa*^DM^ AML as well as in *CEBPA*^-^*TET2* co-mutated patients. This is accompanied by gain of *Gata2* promoter DNA methylation. These findings are consistent with previous data showing that *Gata2* expression is TET2-dependent, as *Gata2* was downregulated in various *Tet2* knockout settings and that forced expression of *Gata2* decreased the competitiveness of both normal and malignant TET2-deficient cells^27, 41–43^. Further paralleling our data, TET2 deficiency in the context of *Flt3*^ITD^ AML has been shown to accelerate leukemia by hypermethylation and consequent silencing of the *Gata2* locus^41^.

Strikingly, we found that while moderate reduction of *Gata2* expression increased competitiveness in *Cebpa*^DM^ AML both *in vivo* and *in vitro*, leukemia cells remain critically dependent on residual GATA2 function. Indeed, homozygous *Gata2* lesions induced a strong inhibitory effect on *Cebpa*^DM^ AML *in vitro*^35^, which was also observed in other AML subtypes as well as in normal hematopoietic stem cells^34, 44–46^. These findings are corroborated by a substantial body of genetic evidence supporting the importance of GATA2 regulation in *CEBPA*-mutant AML. First, heterozygous *GATA2* lesions frequently co-occur with *CEBPA*^DM^ ^4, 8, 16, 17, 30–32, 47–50^. Secondly, *GATA2* allele-specific expression (ASE) is strongly associated with *CEBPA*^DM^ AML and is neither found in AML with reduced *CEBPA* expression (i.e. t(8;21)) nor in *CEBPA*-silenced AML^51^. Thirdly, *TET2*^MUT^ and *GATA2*^MUT^ rarely co-occur in *CEBPA*^DM^ AML, and *GATA2*^MUT^ are less frequently found in *TET2-*deficient *CEBPA*^DM^ compared to *TET2-*proficient *CEBPA*^DM^ AML. Finally, we showed that mutations in *GATA2* and *TET2* are overrepresented in AML cases with high *CEBPA* expression. This supports the notion that unfavorable, high *GATA2* levels in AML promoted by the CEBPA-TET2 axis are not limited to *CEBPA*^DM^ AML, but also include cases where *CEBPA* expression is high for other reasons. Further, this model also suggests that a major proleukemic effect of TET2 deficiency is to rebalance *GATA2* levels in the context of *CEBPA*^DM^ AML (Illustrated in **Figure 7B**).

*GATA2* expression is mainly driven by the conserved *G2*DHE in normal myeloid progenitors and leukemic blasts by promoting expression from the hematopoietic specific *Gata2 V2* promoter^37, 38, 42, 52^. Our data demonstrate that CEBPA plays a key role in regulating *G2*DHE activity. Specifically, we show that the hypermorphic effects of *CEBPA*^DM^ ^11^, and experimental models thereof, result in increased *GATA2* expression compared to *CEBPA*^WT^, and that CEBPA deficiency resulted in reduced *Gata2* levels. Secondly, we observed increased CEBPA binding to the *G2*DHE in *Cebpa*^DM^ AML compared to normal progenitors and found that deletion or mutagenesis of the CEBPA-bound region of the enhancer resulted in lower expression of *Gata2* in *Cebpa*^DM^ cells. In further support of a role of CEBPA, the *G2*DHE is highly active in *CEBPA*^DM^ AML, with both elevated eRNA expression and levels of H3K27ac^51^. An equally important role for CEBPA is observed in the context of inv(3) and t(3;3) AML in which inversions, translocations, and rearrangements involving the *EVI1* gene at the *MECOM locus,* lead to hijacking of the *G2*DHE to promote *EVII* expression at the expense of *GATA2* expression thus resulting in *GATA2* haploinsufficiency^53–56^. Here, *EVI1* expression was found to be downregulated following knockdown of *CEBPA* in inv(3) AML cells, and mutation of the CEBPA binding site in the hijacked enhancer reduced enhancer activity^57^. In this context, *CEBPA*^MUT^ would not be favorable, and these lesions are indeed underrepresented in inv(3) and t(3;3) AML^58–60^.

We hypothesize that CEBPA recruits TET2 and thus mediates DNA demethylation of the *Gata2 V2* promoter in a CEBPA- and TET2-dependent manner. Indeed, *Gata2 V2* levels were decreased, and *Gata2 V2* promoter DNA methylation was increased upon *Cebpa* depletion in a leukemic setting where CEBPA is dispensable for maintenance of the leukemia. The concept of CEBPA as a recruiting factor for TET2 is supported by previous findings showing that both the p30 and p42 isoforms of CEBPA interact with TET2 via the DNA binding domain of CEBPA^61, 62^. Further, CEBPA binds preferentially to methylated DNA^61, 63^, and has been classified as a binding site-directed DNA demethylation-inducing transcription factor^61, 64^. Interestingly, TET2 binds genomic regions that are enriched for CEBP-motifs in myeloid cells, particularly in myeloid enhancers such as the *G2*DHE^26, 61^. Moreover, knockdown or knockout of *Tet2* leads to impaired upregulation of myeloid-specific genes upon *Cebpa* induction, with corresponding increased promoter methylation^65^. Also, in *TET2*^MUT^ or *Tet2*^−/−^ leukemia an enrichment of CEBP-motifs at or near hypermethylated CpGs was observed^26, 66^. Importantly, AML with silenced *CEBPA* is associated with DNA hypermethylation, a feature that is not present in *CEBPA*^DM^ AML, which may suggest a broader function of CEBPA in recruitment of TET2^67^. Given the well-known issues with current TET2 antibodies, we have been unable to ChIP TET2 in our cells, despite numerous attempts^68^. Despite this caveat, we conclude that a growing body of evidence is supporting an important role of CEBPA in recruitment of TET2 to DNA, promoting demethylation and transcription of target genes.

While our findings suggest that *GATA2*^MUT^ and *TET2*^MUT^ both converge at rebalancing the increased expression of *GATA2* in *CEBPA*^DM^ AML, patients with *CEBPA*^DM^ and *GATA2*^MUT^ have a more favorable prognosis^16, 30–32, 47^ than patients harboring the *CEBPA*^DM^ and *TET2*^MUT^ combination^16, 17^. This suggests that while GATA2 deregulation plays an important role in leukemogenesis in the *CEBPA*^MUT^ context, *TET2* deficiency may likely contribute to malignancy through additional mechanisms that shall remain subject of another study. Of clinical interest, we find that TET2 deficiency renders *Cebpa*^DM^ AML sensitive to 5*-*AZA and that TET2-deficient cells lose their proliferative advantage over TET2-proficient cells following 5-AZA treatment. In agreement with TET2-dependent *Gata2* expression, our and previous results show that 5-AZA treatment derepresses *Gata2* expression in TET2-deficient cells^42^. Intriguingly, *CEBPA*^CT^ mutations have recently been reported to sensitize AML to treatment with hypomethylating agents by disrupting the inhibitory interaction with DNMT3A mediated by the wild-type CEBPA bZIP domain^69^. Taken together, this suggests that demethylating agents could be a particularly feasible treatment option in *CEBPA*^DM^*TET2*^MUT^ patients.

In conclusion, our results reveal that *GATA2* is a conserved target of the *CEBPA-TET2* mutational axis in *CEBPA*^DM^ AML and we propose an intricate mechanism by which elevated CEBPA p30 levels mediate recruitment of TET2 to regulatory regions of the *Gata2* gene to promote its expression. We demonstrate that increased GATA2 levels are disadvantageous to *CEBPA*^DM^ leukemic cells and that this can be counteracted by TET2 loss thus providing an explanation for the co-occurrence of CEBPA and TET2 lesions in AML. Finally, increased *Gata2* promoter methylation, inflicted by TET2 deficiency, can be restored by demethylating 5-AZA treatment, thereby providing entry points for the development of rational targeted therapies in AML patients with these mutations.

## ACKNOWLEDGEMENT

Work in the Porse lab was supported by grants from the Copenhagen University Hospital, the Capital Region of Copenhagen and through a center grant from the Novo Nordisk Foundation (Novo Nordisk Foundation Center for Stem Cell Biology, DanStem; Grant Number NNF17CC0027852). This work has been performed in the context of the Danish Research Center for Precision Medicine in Blood Cancers funded by the Danish Cancer Society (R223-A13071) and Greater Copenhagen Health Science Partners.

Work in the Grebien lab was supported by European Union’s Horizon 2020 research and innovation program (European Research Council grant agreement No 636855 and Austrian Science Fund (FWF), grants no. TAI-490 and P35628. ASW was supported by grants from The Lundbeck Foundation (R303-2018-2868) and The Swedish Research Council (2015-00517). EH was supported by a grant from Forschungsförderung from the Fellinger Krebsforschungsverein.

We thank members of the Grebien and Porse laboratories for discussions. We thank Anna Fossum for excellent research assistance.

The authors declare no potential conflicts of interest.

## AUTHOR CONTRIBUTIONS

EH, ASW, GM, MBS, ER, TDA, and CG performed experiments. EH, ASW, AW, TE, MBS, and SP analyzed data. EH, ASW, FG, and BTP designed experiments. JF contributed essential material. MM and TH provided clinical data. EH, ASW, FG, and BTP drafted the manuscript. All authors have proofread and approved the final version of the manuscript.

## MATERIALS AND METHODS

### Patient data

#### Assessment of mutational status

To evaluate co-occurring mutations in *CEBPA*^DM^ AML cases, data from published studies^3–8, 17^ including >40 *CEBPA*^DM^ cases were extracted, and co-occurring mutations were evaluated (**Supplemental table 1**). To determine frequencies of target gene mutations between *CEBPA*^DM^ AML cases with *TET2*^MUT^ compared to TET2 wild-type (*TET2*^WT^) AML cases, data from published studies^3–5, 7, 8, 17, 49^ with specified mutational status including >40 *CEBPA*^DM^ cases or corresponding cohorts were extracted and co-occurring mutations in *TET2*, *GATA2, WT1, CSF3R,* and *ASXL1* were evaluated (**Supplemental table 2A–E**). To examine how the mutational status of *TET2* and *GATA2* were affected by *CEBPA* expression levels in AML, we utilized the publicly available data from the Beat AML cohort (Oregon Health & Science University; OHSU)^1^, including 382 cases for which mutation and mRNA expression data were available. The cases were stratified based on *CEBPA* mRNA expression levels (z-score ±1.0 relative to all samples; *CEBPA*^HIGH^ n=45 and *CEBPA*^LOW^ n=61) and frequencies of *CEBPA*, *TET2* and *GATA2* mutations were determined.

#### Survival analysis

The clinical data set comprises 298 patients with *CEBPA* mutations (MLL Münchner Leukämielabor GmbH), of which 154 harbored biallelic *CEBPA* mutations. Out of these 119 had specified TET2 mutational status and were included in the analyses (*CEBPA*^DM^*TET2*^WT^ n=84, *CEBPA*^DM^*TET2*^MUT^ n=35).

#### Gene expression

The Beat AML dataset used in this study is available at http://vizome.org/aml and comprises 25 patients with *CEBPA* mutations (CEBPA^NT^ and/or *CEBPA*^CT^) for which mutation and mRNA expression data is available. For the gene expression analysis, we excluded patients, which had co-occurring mutation(s) in *WT1* or *IDH1/2* since these have been shown to interfere with TET2 function^70–73^ as well as two patients with low *CEBPA* variant allele frequency (VAF). Gene expression analysis was conducted on data from 16 *CEBPA*-mutant patients of which 5 have a co-occurring mutation in *TET2* (*TET2*^MUT^) (**Supplemental table 3**). Differential expression analysis was performed with DESeq2^74^ (v. 1.26.0) and default parameters.

### *In vitro* experiments

#### Competitive CRISPR-targeting

For generation of *Tet2* or *Gata2* mutated clones, *Cebpa*^p30/p30^ cells^35^ were electroporated with ribonucleoparticles containing recombinant Cas9 nuclease from Streptococcus pyogenes (Sp) (#1081058, IDT), tracrRNA (#1075927, IDT) and crRNAs (Alt-R® CRISPR-Cas9 crRNA, IDT) targeting *Tet2* and *Gata2*, respectively. crRNAs were designed using the CHOPCHOP^75^ web tool (chopchop.cbu.uib.no) (**Supplemental table 4**). crRNA and tracrRNA molecules were complexed at room temperature and assembled with recombinant SpCas9 according to manufacturer’s protocols (IDT). Pools of *Tet2-* or *Gata2*-targeted cells were screened at regular intervals to monitor outgrowth of subpopulations. The genomic regions that were targeted with CRISPR/Cas9 technology were PCR-amplified, Sanger sequenced and analyzed with the online tool Tracking of Indels by DEcomposition (TIDE)^76^ for insertions or deletions (indels) in the targeted region. Primers for PCR are provided in **Supplemental table 5**.

#### Gata2 enhancer CRISPR-targeting

sgRNA sequences targeting the *Gata2* distal hematopoietic enhancer (G2DHE) were obtained from the UCSC Genome Browser^77^ (genome.ucsc.edu) and targets with a high predicted cleavage (Doench/Fusi 2016 Efficiency > 55) selected (**Supplemental table 4**). *Cebpa*^p30/p30^ cells were co-transduced with pLenti-hU6-sgG2DHE_A/B-IT-PGK-iRFP and LentiGuide-sgG2DHE1-11-Puro-IRES-GFP. GFP^+^iRFP670^+^ cells were sorted via fluorescence-activated cell sorting (FACS) and frozen for subsequent analysis. For Figure 5E, *Cebpa*^p30/p30^ were transduced with LentiGuide-sgG2DHE-Puro-IRES-GFP, followed by FACS for GFP+ cells and subsequent analysis.

### *In vivo* experiments

Experiments were carried out according to protocols approved by the Danish Animal Ethical Committee. Mice were bred and housed locally at the Department of Experimental Medicine at the University of Copenhagen. The mice were housed in a temperature- and humidity-controlled room with a 06:00–18:00 h light cycle and fed a standard chow diet and tap water *ad libitum*. We used *Tet2*^fl/fl^ ^78^, *Cebpa*^p30/+^ ^13^, *Cebpa*^+/fl^ ^79^ and *Mx1-Cre*^+^ ^80^ lines to generate *Tet2*-deficient and *Cebpa*-mutant compound lines. The following genotypes were used for experiments: *Cebpa*^fl/p30^*Tet2*^+/+^, *Cebpa*^+/fl^*Tet2*^fl/fl^, *Cebpa*^fl/p30^*Tet2*^fl/fl^, *Cebpa*^fl/p30^*Tet2*^+/+^*Mx1-Cre*^+^, *Cebpa*^+/fl^*Tet2*^fl/fl^*Mx1-Cre*^+^, and *Cebpa*^fl/p30^*Tet2*^fl/fl^*Mx1-Cre*^+^. We used *iMLL-AF9*^+^ ^81^, *Cebpa*^+/fl79^ and *R26-CreER*^+^ ^82^ lines to generate an *iMLL-AF9*^+^*Cebpa*^fl/fl^*R26-CreER*^+^ compound line. Primers for genotyping in are provided in **Supplemental table 5**.

#### Leukemia initiation

C57BL6/6J.SJL congenic recipients (female, 10–12 weeks old) were lethally irradiated (900 cGy) 12–24h prior to intravenous injection with 1×10^6^ bone marrow (BM) cells from individual donor mice. The mice were given Ciprofloxacin (100 mg/l in acidified water; #17850 Sigma-Aldrich) in the drinking water to prevent infections 3 weeks post-irradiation. Recipient mice were allowed to recover for 6 weeks post-transplantation before Cre-LoxP recombination was induced by two intraperitoneal injections of Poly(I)-Poly(C) (300 μg in 200 μl PBS; #27-4732-01 GE Healthcare) with 48 h rest in-between. The day of the first injection was set as time-point zero for the survival study and mice were monitored for leukemia development and euthanized when moribund. To follow leukemia initiation in the recipients, a subgroup of mice was subjected to blood and BM sampling at 12, 24, and 36-week time-points. BM from moribund mice was collected and frozen (10% DMSO in FBS; #D8418 Sigma-Aldrich, #HYCLSV30160.03 Hyclone) for subsequent FACS and analysis.

#### Competitive shRNA-knockdown

C57BL6/6J.SJL recipients (female, 10–12 weeks old) were sub-lethally irradiated (500 cGy) 12–24h prior to being intravenously injected with a 1:1 mix of *Cebpa*^p30/p30^ cells^13^ transduced with shRNA targeting *Gata2* (**Supplemental table 6**) or with control-shRNA as previously described^83^. The ratio of *Gata2*- or control-shRNA-GFP^+^ to control-shRNA-YFP^+^ cells was analyzed by flow cytometry four weeks later.

#### 5-azacytidine treatment

C57BL6/6J.SJL recipients (female, 10–12 weeks old) were sub-lethally irradiated (500 cGy) 12–24h prior to being intravenously injected with 1×10^5^ thawed live BM cells from moribund secondary recipient mice. The mice were given Ciprofloxacin in the drinking water to prevent infections 3 weeks post-irradiation. The mice received intraperitoneal injections with the demethylating agent 5-azacytidine (2.5 mg/kg/day in saline; #A2385 Sigma-Aldrich) at days 6–10 and 20–24 post-transplantation. The time of the BM cell injection was set as time-point zero for the survival study and mice were monitored and euthanized when moribund. To evaluate the effects of short-term 5-azacytidine treatment, recipient mice were treated at days 13-15 and euthanized 24 hours after the last injection. BM was collected for FACS, and sorted cells were frozen for subsequent analysis.

### Immuno-staining

Detailed in **Supplemental Material and Methods**.

### *Ex vivo* cell culture

#### Establishment of ex vivo iMLL-AF9^+^Cebpa^fl/fl^R26-CreER^+^ lines

Sorted GMPs from *iMLL-AF9*^+^*Cebpa*^fl/fl^*R26CreER*^+^ mice, were cultured in MethoCult (M3434; #03434, Stemcell technologies) supplemented with doxycycline (1 μg/ml; #D9891 Sigma-Aldrich) for three replatings to induce expression of the MLL-fusion protein.

#### Cebpa knockout

Leukemic *iMLL-AF9*^+^*Cebpa*^fl/fl^*R26CreER*^+^ cells were cultured in RPMI 1640 medium (#21875034, Gibco) supplemented with FBS (10%), Penicillin-Streptomycin (1%), doxycycline (1 μg/ml), and cytokines m-IL-3 (6 ng/ml), m-SCF (50 ng/ml), and h-IL-6 (10 ng/ml). After two days, 4-hydroxytamoxifen (4-OHT; 1 μM; #H7904 Sigma) or vehicle was added to the cell culture medium to activate *Cre-LoxP* recombination. Three days later cells were isolated and either frozen or resuspended in RA1 buffer (NucleoSpin RNA XS, # 740902 Macherey-Nagel). The experiment was run in triplicates for treatment and vehicle, with two biological replicates.

### High-throughput sequencing and bioinformatic analyses

Detailed in **Supplemental Material and Methods**.

### ShRNA knockdown of *Gata2*

#### Cloning of shRNA into pMLS vector

Murine shRNAs targeting *Gata2* (sh*Gata2*) were cloned into MSCV-LTRmir30-SV40-GFP vector. Targeting sequences were identified from the Mission® shRNA library (**Supplemental table 6**) and the sense and anti-sense sequences were incorporated with a miR-30-loop to generate a 97-mer target sequence. Oligonucleotides were amplified by PCR using miR30 common primers (**Supplemental table 5**), which include restriction sites for *XhoI* and *EcoRI*. The resulting 138-mer PCR amplicons and the vector were digested with *XhoI* and *EcoRI* and products were ligated using T4 DNA Ligase (#15224025 Invitrogen). Bacterial transformation was performed to amplify individual ligation products, and correct inserts were verified by Sanger Sequencing. These, together with vectors containing a control non-targeting sequence (MSCV-LTRmir30-SV40-GFP and MSCV-LTRmir30-SV40-YFP), were used in subsequent transfection/transduction experiments, as previously described^83, 84^.

#### Transduction of Cebpa^p^^30/p^^30^ cells

Retroviral transduction was done as previously described^83^. Briefly, retroviral supernatants were generated by transfection of Phoenix E cells. For transduction, retroviral supernatant was added onto retronectin-coated (1:25; #T100B TaKaRa) non-tissue culture treated plates and centrifuged at 2000 ×g for 60 min at 4 °C. After aspiration of the supernatant, *Cebpa*^p30/p30^ cells were seeded at a density of 0.5–1×10^5^ cells/cm^2^. The transduction was repeated the following day, and the cells were cultured for 24 h prior to FACS sorting of transduced (GFP^+^/YFP^+^) cells on a BD FACSAria^TM^ III (BD Bioscience). The efficiency of shRNA-mediated gene expression knockdown was assessed with qPCR and cells were used for transplantation and assessment of their competitiveness *in vivo*.

### Quantitative PCR

RNA from sorted blasts or *ex vivo*-cultured cells was isolated using NucleoSpin RNA XS kit (#740902 Macherey-Nagel) or RNeasy Mini Kit (#74104 Qiagen) according to the manufacturers’ instructions and converted to cDNA using ProtoScript First Strand cDNA Synthesis Kit (#E6300 New England BioLabs). Quantitative PCR (qPCR) to assess knockdown efficiency was run using TaqMan Fast Advanced Master Mix (#4444556 Applied Biosystems^TM^) and TaqMan assay for *Gata2* (Mm00492301_m1 FAM-MGB), in duplex with housekeeping gene *18S* (Hs99999901_s1 VIC-MGB-PL). qPCR to evaluate mRNA levels of total *Gata2*, *variant 1* (*V1*) and *variant 2* (*V2*), respectively, was run in duplex using LightCycler 480 SYBR Green I Master (#04887352001 Roche) with primers for *Gata2* and housekeeping gene *Actb* and *Gapdh*^39^ (**Supplemental table 5**). Gene expression was calculated with the 2^−ΔΔct^ method.

RNA from *Cebpa*^p30/p30^ cell lines was isolated using RNeasy Plus Mini Kit (#74134 Qiagen) according to the manufacturer’s instructions and converted to cDNA with RevertAid First Strand cDNA Synthesis Kit (K1622). qPCR was run using SsoAdvanced Univ SYBR Grn Suprmix (#1725271, Bio-Rad Laboratories Ges.m.b.H.) and primers for *Gata2* and *Gapdh* (**Supplemental table 5**).

### Bisulfite PCR

DNA was isolated using DNeasy Blood and tissue kit (#69504 Qiagen) and the DNA was bisulfite converted using EZ-DNA Methylation Gold Kit (#D5005 Zymo Research), both according to the manufacturer’s instructions. PCR was run using Pfu Turbo Cx Hotstart DNA polymerase (#600410 Agilent) with primers targeting a part of the CpG island in the *Gata2 V2* promoter region (**Supplemental table 5**). After verification of their correct size, PCR products were cloned using Zero Blunt Topo PCR Cloning kit (#450245 Invitrogen) and single colonies were picked and amplified. Plasmid DNA was isolated using NucleoSpin Plasmid EasyPure (#740727.250 Macherey-Nagel), the correct insert size was verified after cleavage with restriction enzyme EcoRI (#R0101 New England Biolabs) and sent for Sanger sequencing using the M13 primer provided with the cloning kit.

### Statistics

Data were analyzed for significance using parametric tests, with prior log-transformation if necessary to achieve normal distribution. Normality was evaluated by Shapiro-Wilk test. Two-group analyses were done using unpaired two-tailed t-test. Multiple-group analyses were done with one-way-ANOVA followed by multiple comparisons correction using Dunnett when comparing to a reference group, or two-way-ANOVA followed by multiple comparisons correction using Šídák test when comparing two independent factors across four groups. Data sets that did not pass normality tests were analyzed by Kruskal-Wallis test followed by multiple comparisons correction using Dunn’s test. Survival curves were analyzed using Mantel-Cox Log-rank test. To compare distributions Wilson/Brown binominal test was used. P-values <0.05 were considered statistically significant. Data was analyzed using GraphPad Prism (v. 9). Data is shown as mean±SEM unless otherwise stated.

### Data availability

The data generated in this study is publicly available in Gene Expression Omnibus (GEO) under accession numbers GSE214224 (RNA-seq and ATAC-seq *in vitro*) and GSE213864 (RNA-seq and WGBS *in vivo*), and within the article and its supplementary files. CEBPA and H3K27Ac ChIP-seq from myeloid progenitor cell model for p30-driven AML is available under GSE158727 (Heyes et al.^12^). CEBPA ChIP from mouse *Cebpa*^+/+^ or *Cebpa*^p30/p30^ GMPs is available under GSE118963 (Jakobsen et al.^11^). TET2 ChIP-seq is available under GSE115972 (Rasmussen et al.^26^). Patient data analyzed in this study were from the Beat AML study (Tyner et al.; accessed through cBioPortal^85^ or Vizome^1^) or from published cohort studies (**Supplemental tables 1 and 2A-E**).

Illustrations were created with BioRender.com

