## Supplement for "*TET2* lesions enhance the aggressiveness of *CEBPA-*mutant AML by rebalancing *GATA2* expression"

### SUPPLEMENTAL MATERIALS AND METHODS

#### Western blotting

Western blotting for TET2 was performed according to standard laboratory protocols, using the following antibodies: anti-TET2 (Santa Cruz, sc-398535) and anti-HSC70 (Santa Cruz, sc-7298).

#### *In vivo* experiments

*Leukemia propagation:* C57BL6/6J.SJL recipients (female, 10–12 weeks old) were lethally irradiated (900 cGy) 12–24h prior to being intravenously injected with $2\times10^5$ thawed live BM cells from moribund donor mice together with $4\times10^5$ freshly isolated BM cells from C57BL6/6J.SJL mice. The day of the injection was set as time-point zero for the survival study and mice were monitored and euthanized when moribund. The mice were given Ciprofloxacin in the drinking water to prevent infections 3 weeks post-irradiation.

#### Immuno-staining

*Flow cytometry:* To analyze the composition of either freshly isolated or thawed cryopreserved BM and blood, cells were stained with fluorescently labelled antibodies. For blood analysis, 50 µl blood was collected from the facial vein and erythrocytes were lysed with lysing buffer (BD Pharm Lyse™, #555899 BD Bioscience). For BM analysis, cells were collected by crushing tibia, femur, and ilium and filtered through a 50 µm filcon cup (#340630 BD Bioscience). Blood or BM cells were washed in PBS with 3% FBS and stained with fluorescently labelled antibodies for 30 min at 4°C (**Supplemental table 7**). For cryopreserved cells, the cells were counterstained with DAPI (1:10000; #D3571 Invitrogen) to separate out dead cells. Fluorochrome-minus-one was used as controls. Flow cytometry data was obtained using a BD FACSAria™ III or a BD LSR II™ (BD Bioscience) and analyzed using FlowJo software (v9; BD Bioscience).

For downstream transcriptional and epigenetic analyses, live donor-derived non-lymphoid and non-erythroid cells ($DAPI^{-}CD45.2^{+}CD3^{-}B220^{-}Ter119^{-}$) were sorted using a BD FACSAria™ III, spun down and cell pellets were either snap-frozen or resuspended in RLT buffer (RNeasy Mini Kit, #74104 Qiagen).

For *ex vivo* cell culture of *iMLL-AF9*$^{+}$*Cebpa*$^{fl/fl}$*R26-CreER*$^{+}$ cells, c-kit$^{+}$ BM cells were enriched by magnetic sorting (mouse CD117 MicroBeads; #130-091-224, Miltenyi Biotec), and granulocyte/monocyte progenitors (GMPs; $Lin^{-}C\text{-}kit^{+}Sca1^{-}CD41^{-}FcgRII^{+}$) were sorted using a BD FACSAria™ III.

*Immunohistochemistry:* To evaluate proliferative status of leukemia cells, cells from BM of moribund mice were spun on glass slides, air-dried, and fixed with methanol (#VWRC20846.292 VWR). After blocking of endogenous peroxidase activity with hydrogen peroxide (1%), slides were stained over night at 4°C with anti-Ki67 antibody (1:50; #ab16667 Abcam) in antibody diluent (S3022 Dako). To visualize the primary antibody, EnVision HRP Rabbit (K4003 Dako) together with Vina Green™ Chromogen Kit (BRR807 Biocare Medical) was utilized according to manufacturer's instructions. The cells were counterstained with Mayer Hematoxylin (#51275 Sigma-Aldrich), dehydrated and coverslips mounted with Entellan (#107960 Sigma-Aldrich). Images were captured using a Leica microscope at 20X magnification and Ki67$^{+}$ cells were quantified out of one hundred cells.

#### *Ex vivo* cell culture

*Establishment of ex vivo Cebpa*$^{-/p30}$*Tet2*$^{+/+}$ *and Cebpa*$^{-/p30}$*Tet2*$^{-/-}$ *lines:* Thawed cryo-preserved cells from primary AML were cultured in Lonza X-Vivo™ 15 cell medium (#BE02-060Q Thermo Fisher Scientific) supplemented with Bovine Serum Albumin in Iscove's MDM (10%; #09300 Stemcell™ Technologies), Penicillin-Streptomycin (1%; #15140122 Gibco), β-mercaptoethanol (0.1 mM; #31350010 Gibco), L-glutamine (2 mM; #25030149 Gibco), and cytokines h-IL-6 (50 ng/ml; #130-093-032 Miltenyi Biotec), m-SCF (50 ng/µl; #250-03 Peprotech), m-IL-3 (10 ng/ml; #213-13 Peprotech), and m-GM-CSF (10 ng/ml; #315-03 Peprotech). Two clones of each genotype (*Cebpa*$^{-/p30}$*Tet2*$^{+/+}$ and *Cebpa*$^{-/p30}$*Tet2*$^{-/-}$) continued to expand beyond 40 days and withstood freeze-thawing, and these clones have been used for further experiments.

*Vitamin C treatment:* Cells were seeded at a density of $2\times10^5$ cells/ml and the cell culture medium was supplemented with vitamin C (100 µg/ml; L-ascorbic acid, #A8960 Sigma Aldrich). Live cells were counted using Solution 13 (AO-DAPI; #910-3013 Chemometec) on a NucleoCounter® NC-3000™ and reseeded at $2\times10^5$ cells/ml every third day. The experiment was run in triplicates for treatment and vehicle, with two biological replicates for each genotype.

*5-azacytidine treatment:* Cells were seeded at a density of $2\times10^5$ cells/ml and medium supplemented with 5-azacytidine (5-AZA; 1 µg/ml; #A2385 Sigma-Aldrich). Live cells were counted using Solution 13 on a NucleoCounter® NC-3000™ and reseeded at $2\times10^5$ cells/ml day three and six. 24 hours later, up to $1\times10^5$ cells were isolated and resuspended in RA1 buffer (NucleoSpin RNA XS, # 740902 Macherey-Nagel). The experiment was run in triplicates for treatment and vehicle, with two biological replicates for each genotype.

#### High-throughput sequencing and bioinformatic analyses

*RNA-sequencing (RNA-seq) of cell line models:* RNA was isolated from $1\times10^6$ cells using RNeasy Plus Mini Kit (#74134 Qiagen) according to the manufacturer's instructions and quality was assessed on a Bioanalyzer 2100 G2939A (Agilent). 1 µg of RNA was used to generate sequencing libraries using QuantSeq 3′ mRNA-Seq Library Prep Kit (FWD) for Illumina, 96 preps (#015.96, Lexogen) and the PCR Add-on Kit for Illumina, 96 rxn (#020.96, Lexogen). The libraries were quantified on a Bioanalyzer 2100 G2939 (Agilent) and pooled in equimolar amounts. Multiplexed libraries were sequenced on a HiSeq4A (Illumina).

*RNA-seq of leukemic cells from in vivo models:* RNA was isolated from $5\times10^5$ sorted cells using RNeasy Mini Kit (#74104 Qiagen) according to the manufacturer's instructions and quality was assessed by RNA 6000 Pico Kit (#5067-1513 Agilent) on a Bioanalyzer 2100 (Agilent). 200 ng RNA was used to generate sequencing libraries using TruSeq RNA Library Prep Kit v2 (#RS-122-2001 Illumina). The libraries were quantified using Qubit dsDNA BR Assay Kit (#32853 Thermo Fisher Scientific) and DNA 1000 Kit (#5067-1504 Agilent) and pooled in equimolar amounts. Multiplexed libraries were sequenced on a NextSeq 500 (Illumina) using NextSeq 500 High Output v2 Kit (75 cycles; #FC-404-2005 Illumina).

*Bioinformatics analyses of RNA-seq data*: RNA-seq analysis for *in vitro Cebpa*$^{p30/p30}$ cells was performed as previously described[1,2]. Quality check was done with FastQC[3] (v. 0.11.4) and preprocessing with PRINSEQ-lite[4], using parameters: -min_len 30 -min_qual_mean 30 -ns_max_n 5 -trim_tail_right 8 -trim_tail_left 8 -trim_qual_right 30 -trim_qual_left 30 -trim_qual_window 5. Remaining reads were aligned against the mouse reference genome (mm10) with BWA[5]. RNA-seq analysis for *in vivo Cebpa*$^{-/p30}$ cells was performed as follows. RNA-seq reads were processed with the bcbio RNA-seq pipeline (https://github.com/bcbio/bcbio-nextgen) and the bcbioRNASeq R package (https://github.com/hbc/bcbioRNASeq). In brief, transcript abundance estimates were obtained using Salmon[6] (v. 0.12.0) against reference transcriptome GRCm38/mm10 ENSEMBL release 94, summarized to gene level and imported into R using tximport[7] (v. 1.10.1). Differential gene expression analysis between the *Cebpa*$^{-/p30}$*Tet2*$^{+/+}$ and *Cebpa*$^{-/p30}$*Tet2*$^{-/-}$ genotype was performed using DESeq2 with standard parameters[8] (v. 1.22.2).

*Gene set enrichment analysis (GSEA):* GSEA was performed using the GSEA software[9,10] (v. 4.1.0).

*Assay for transposase-accessible chromatin-sequencing (ATAC-seq):* ATAC-seq was performed as previously described[2].

*Bioinformatics analyses of ATAC-seq data:* Analysis of ATAC-seq was performed as previously described[2]. HOMER[11] (v. 4.11) was used to identify motifs enriched in the ATAC peaks.

*Bisulfite whole genome sequencing (WGBS):* DNA was isolated from $1\times10^6$ sorted cells using DNeasy Blood and tissue kit (#69504 Qiagen) according to the manufacturer's instructions. Bisulfite conversion of DNA was done according to manufacturers' instructions using EZ-DNA Methylation Gold Kit (#D5005 Zymo Research). Quality of bisulfite treated DNA was assessed by RNA 6000 Pico Kit (#5067-1513 Agilent) on a Bioanalyzer 2100. Libraries of bisulfite converted DNA were prepared using Pico Methyl-Seq Library Prep Kit (#D5455 Zymo Research) according to manufacturer's instructions and the final concentration and quality of the libraries was assessed using Qubit dsDNA HS Assay Kit (#Q32854 Thermo Fisher Scientific) and High Sensitivity DNA Analysis Kit (#5067-4626 Agilent) on a Bioanalyzer. Duplexed libraries were sequenced on a NextSeq 500 (Illumina) using NextSeq 500 High Output v2 Kit (75 cycles).

*Bioinformatics analyses of WGBS data:* Reads were trimmed and filtered using Trim Galore[12] (v. 0.4.3) with default parameters, quality was assessed before and after using FastQC[3] (v. 0.11.7). Trimmed reads were aligned to the mouse genome assembly GRCm38 (mm10) using Bismark[13] (v. 0.19.1) with option --non_directional (other parameter left at default values; this used Bowtie2[14] (v. 2.2.8) with -q --score-min L,0,-0.2 --ignore-quals). After deduplication of alignments (using deduplicate_bismark), the methylation information for individual cytosines was extracted using bismark_methylation_extractor (--cytosine_report --comprehensive --gzip). To quantify DNA methylation of gene bodies and promoters (1000 bp up- and downstream of transcription start sites), we used the weighted methylation level (i.e., summarizing over all CpG positions in the given region, the number of reads supporting methylated cytosine divided by the number of all reads covering these positions). Plots of average methylation levels across extended gene bodies were generated using deepTools[15] (v.3.1.3) computeMatrix (scale-regions -m 4000 -a 1000 -b 1000 --unscaled5prime 1000 --unscaled3prime 1000) and plotProfile, for which Bismark-generated bedGraph files were converted to BigWig format (using UCSC's bedGraphToBigWig[16] (v. 4)).

*Bioinformatic analyses of chromatin immunoprecipitation-sequencing:* ChIP-seq data from *in vitro Cebpa*$^{p30/p30}$ cells was processed as described[2]. ChIP-seq data from *in vivo Cebpa*$^{p30/p30}$ cells was processed as follows; raw reads derived from CEBPA (*Cebpa*$^{+/+}$ and *Cebpa*$^{p30/p30}$) ChIP-seq experiments were mapped to mouse (mm10) genome assembly using Bowtie2[14] (v. 2.3.4.3). We used uniquely mapped and PCR duplicates (exact copies) collapsed as one read and extended to their fragment length by determining the read extension size using MACS2[17] (v. 2.1.0.20151222; predicted parameter). Raw read counts were normalized to TPM using deepTools[15] (v. 3.3.1; bamCoverage). Raw read counts (CEBPA binding levels) mapping to *Gata2* promoter and enhancer regions were computed using bedtools[18] (v. 2.30.0; multicov), and the differences in CEBPA binding between *Cebpa*$^{+/+}$ and *Cebpa*$^{p30/p30}$ conditions were computed using DESeq2[8] (v. 1.30.1).

**Chromatin Immunoprecipitation (ChIP)-qPCR**
ChIP for CEBPA was performed as previously described[1]. The sequences used for qPCR are listed in **Supplemental table 5**.

### SUPPLEMENTAL FIGURES

**Supplemental figure 1, related to Figure 1:**

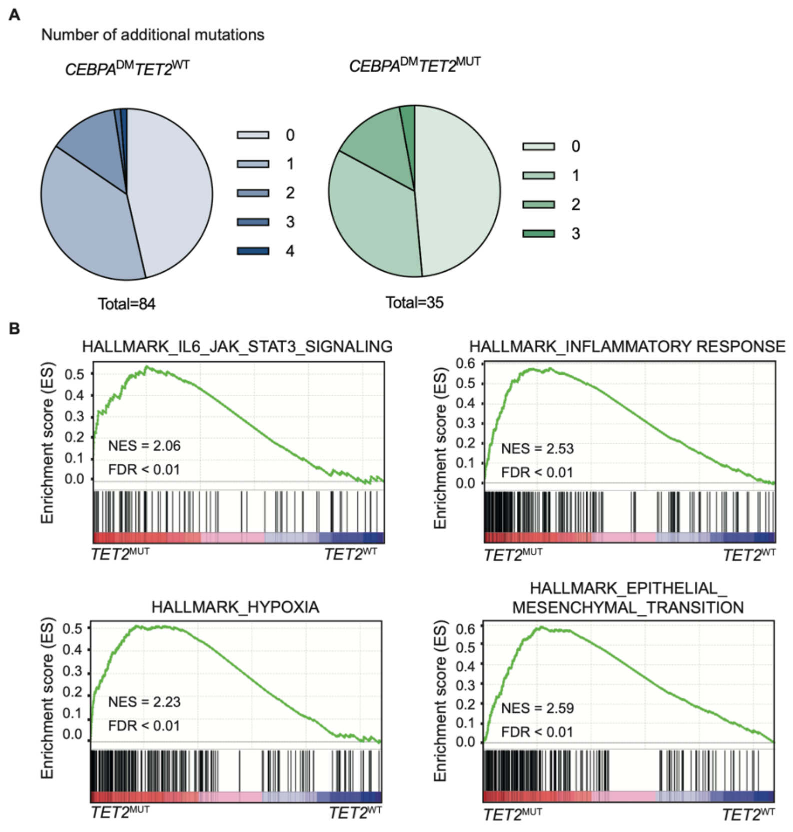


(**A**) Number of additionally accumulated mutations in *CEBPA*$^{DM}$*TET2*$^{WT}$ and *CEBPA*$^{DM}$*TET2*$^{MUT}$ in the patient cohort shown in Figure 1B. (**B**) Gene set enrichment analysis (GSEA) of the *TET2*$^{WT}$ versus *TET2*$^{MUT}$ patients from the cohort of *CEBPA*-mutant patients in the Beat AML dataset.
NES=normalized enrichment score; FDR=false discovery rate

**Supplemental figure 2, related to Figure 2:**

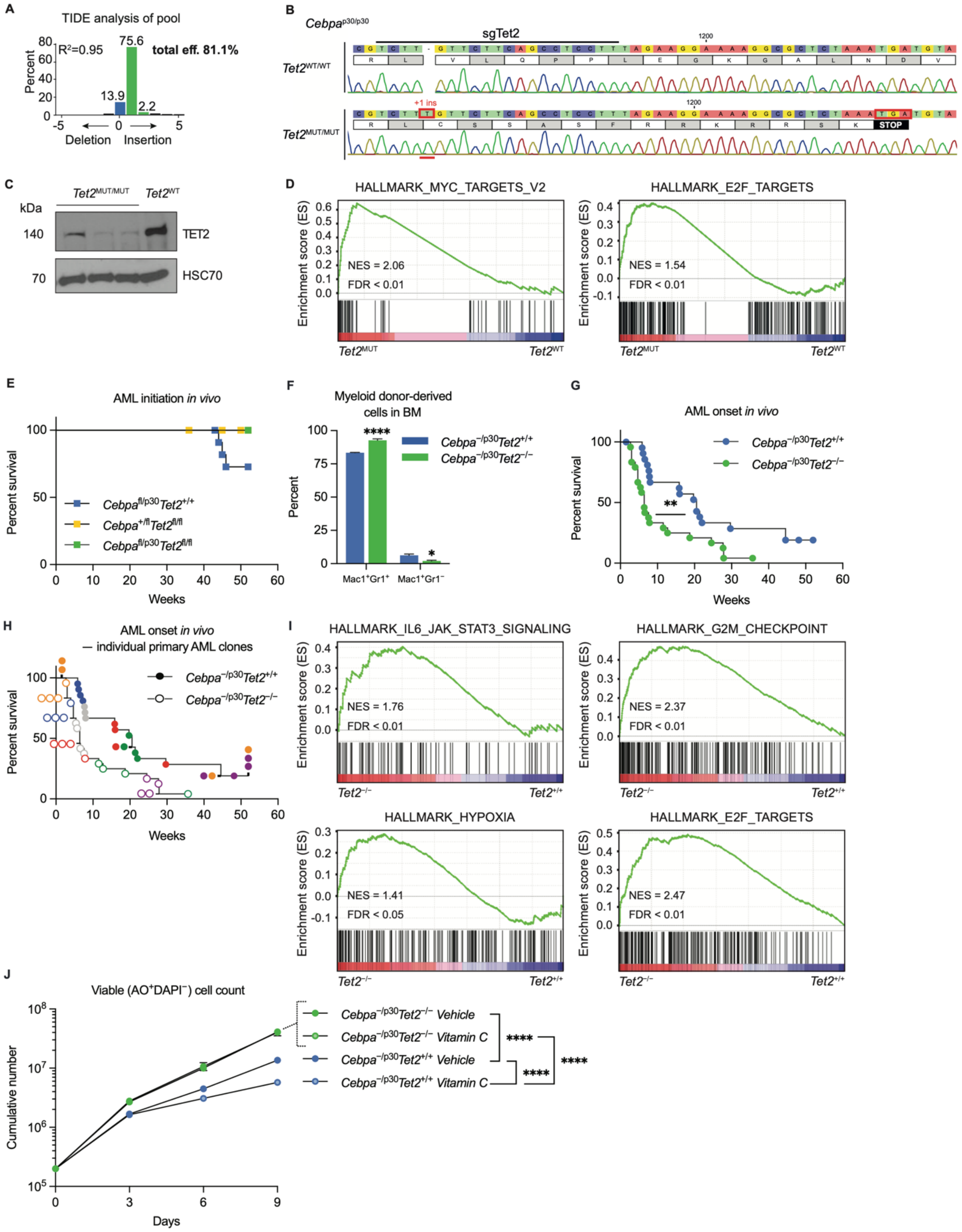


(**A**) Tracking of indels by decomposition (TIDE) of a mixed pool of *Cebpa*$^{p30/p30}$ cells 3 months post CRISPR-targeting of *Tet2*. (**B**) Sanger sequencing analysis of the mutated *Tet2* region in a *Cebpa*$^{p30/p30}$*Tet2*$^{WT/WT}$ and a *Cebpa*$^{p30/p30}$*Tet2*$^{MUT/MUT}$ clone. (**C**) Western blot showing TET2 protein levels in *Cebpa*$^{p30/p30}$*Tet2*$^{MUT}$ clones compared to *Cebpa*$^{p30/p30}$*Tet2*$^{WT}$. (**D**) GSEA plots for selected upregulated gene sets of *Cebpa*$^{p30/p30}$*Tet2*$^{MUT}$ versus *Cebpa*$^{p30/p30}$*Tet2*$^{WT}$ (n=5–7 per group). (**E**) AML initiation after transplantation and Cre-LoxP recombination of

control mice not carrying *Mx1-Cre* (n=7–13/group). (**F**) Myeloid markers in donor-derived cells in bone marrow from moribund mice assessed by flow cytometry (n=3 per group). (**G**) Survival of lethally irradiated secondary recipient mice after transplantation of leukemic BM from moribund primary recipient mice together with normal BM cells (n=23–24/group). (**H**) Survival of secondary recipient mice with primary AML clone indicated by color (Primary AML n=6; secondary recipient per primary AML n=3–4). (**I**) GSEA plots for selected upregulated gene sets of *Cebpa*$^{-/p30}$*Tet2*$^{-/-}$ versus *Cebpa*$^{-/p30}$ *Tet2*$^{+/+}$ (n=3 per group). (**J**) Cell growth *ex vivo* assessed in the presence of Vitamin C or vehicle indicated by viable (AO$^{+}$DAPI$^{-}$) cell count (2 biological and 3 technical replicates per genotype).
*=P<0.05, **=P<0.01, ****=P<0.0001
NES=normalized enrichment score; FDR=false discovery rate

**Supplemental figure 3, related to Figure 3:**

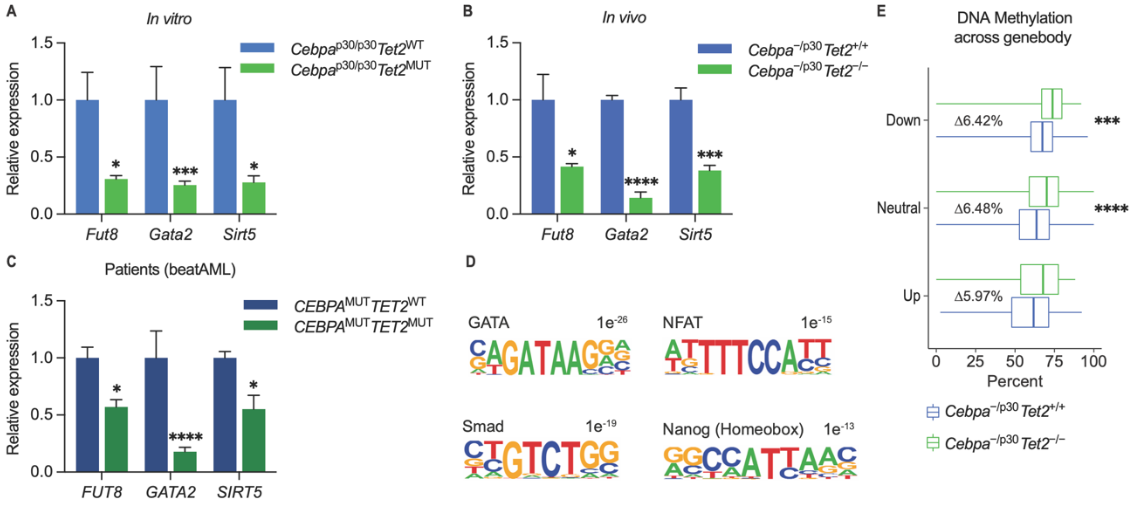


(**A**) Target RNA levels upon TET2-deficiency in *Cebpa*$^{p30/p30}$ cells (n=5–7/group), and (**B**) *Cebpa*$^{-/p30}$ leukemic blasts (n=3/group), as well as (**C**) *CEBPA*$^{MUT}$*TET2*$^{MUT}$ vs. *CEBPA*$^{MUT}$*TET2*$^{WT}$ cases from the Beat AML study [19] (n=5–11/group). Significance of the data is indicated as adjusted P-values. (**D**) Motifs enriched in promoter regions shown in Figure 3C. (**E**) Percent DNA methylation (mC) in sorted AML blasts across the gene bodies of up-, non- vs. down-regulated genes. Boxes indicates the lower quartile-median-upper quartile and whiskers max–min.

*=P<0.05, ***=P<0.001, ****=P<0.0001

**Supplemental figure 4, related to Figure 4:**

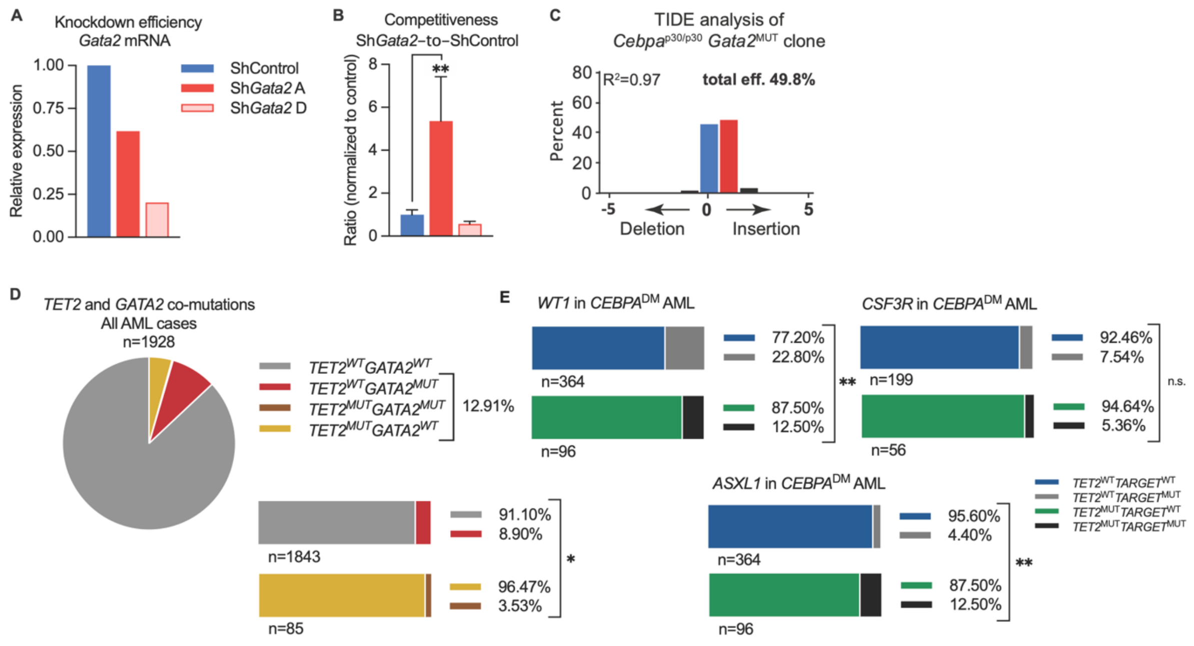


(**A**) *Gata2* mRNA levels in shRNA transduced *Cebpa*$^{p30/p30}$ leukemic cells prior to transplantation and (**B**) *in vivo* Sh*Gata2*-to-ShControl cell ratio normalized to control at 28 days after transplantation of the cells into sub-lethally irradiated recipients (n=4/group). (**C**) TIDE analysis of a representative *Cebpa*$^{p30/p30}$*Gata2*$^{mut}$ clone. **(D)** Presence or absence of *GATA2* mutations (*GATA2*$^{MUT}$) in all AML cases with or without *TET2* mutations (*TET2*$^{MUT}$) (*TET2*$^{MUT}$*GATA2*$^{WT}$ n=82, *TET2*$^{MUT}$*GATA2*$^{MUT}$ n=3, *TET2*$^{WT}$*GATA2*$^{WT}$ n=1679, *TET2*$^{WT}$*GATA2*$^{MUT}$ n=164). **(E)** Presence or absence of *WT1, CSF3R,* and *ASXL1* mutations in *CEBPA*$^{DM}$ AML cases with or without *TET2* mutations.
*=P<0.05, **=P<0.01

**Supplemental figure 5, related to Figure 5:**

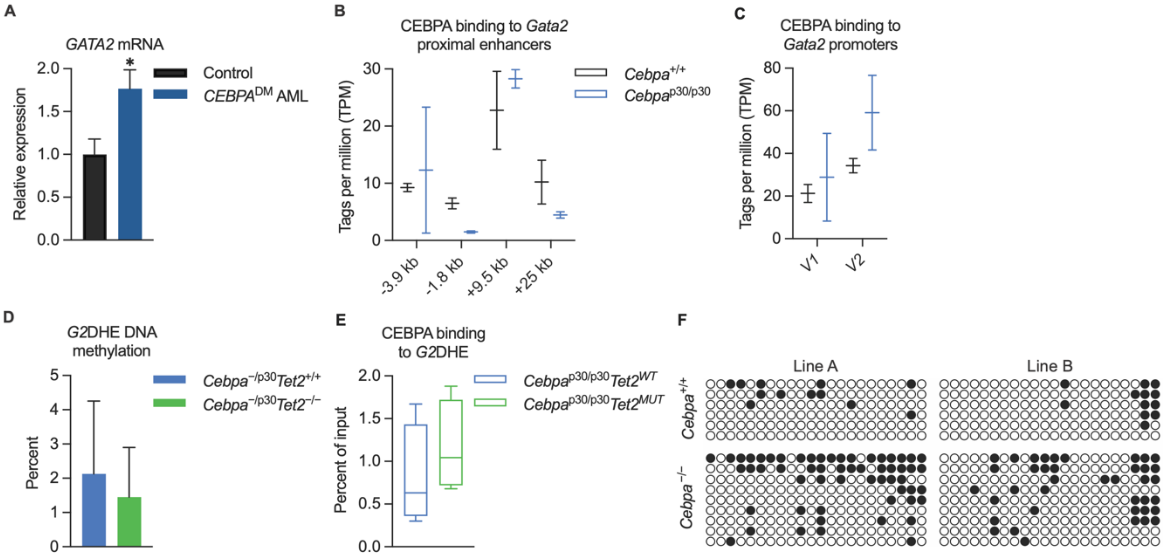


**(A)** *GATA2* mRNA expression in human *CEBPA*$^{DM}$ AML vs. normal hematopoietic cells (n=2–4 per group), data from Jakobsen et al.[20]. (**B**) CEBPA-binding at *Gata2* enhancer, and (**C**) promoter regions in mouse *Cebpa*$^{p30/p30}$ L-GMPs vs. normal GMPs (n=2 per group), data from Jakobsen et al.[20]. (**D**) *G2*DHE DNA methylation in *Cebpa*$^{-/p30}$*Tet2*$^{-/-}$ and *Cebpa*$^{-/p30}$*Tet2*$^{+/+}$ blasts (n=2–3 per group). (**E**) CEBPA binding at *G2*DHE in *Cebpa*$^{p30/p30}$*Tet2*$^{WT}$ and *Cebpa*$^{p30/p30}$*Tet2*$^{MUT}$ cells. (**F**) Schematic overview of CpG methylation in the *Gata2 V2* promoter upon *Cebpa* knockout in two lines of MLL-AF9 leukemia (White circles=C and Black circles=mC) (2 biological replicates per genotype).
*=P<0.05

**Supplemental figure 6, related to Figure 6:**

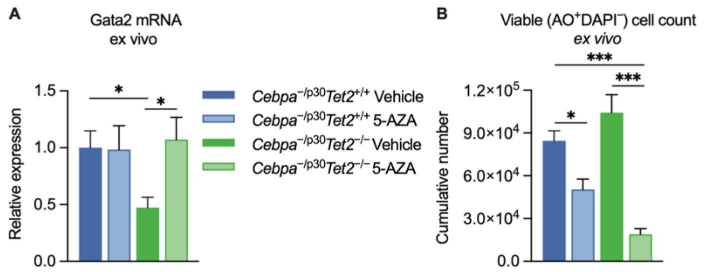


*Ex vivo* culture of leukemic blasts in response to demethylating agent 5-Azacytidine (5-AZA) or vehicle. (**A**) *Gata2* mRNA expression, and (**B**) cell growth, measured as viable ($AO^{+}DAPI^{-}$) cell count, assessed after 6 days in the presence of 5-AZA or vehicle (2 biological and 3 technical replicates per genotype).
*=P<0.05, ***=P<0.001

### SUPPLEMENTAL TABLES

**Supplemental table 1: Co-occurring mutations in *CEBPA*$^{DM}$ AML cases**

| **Cohort** | ***CEBPA*$^{DM}$ (n)** | ***GATA2*** | ***TET2*** | ***WT1*** | ***NRAS*** | ***FLT3*** | ***CFSR3*** | ***DNMT3A*** | ***IDH1/2*** | ***ASXL1*** | ***KIT*** | ***EZH2*** | ***RUNX1*** | ***KRAS*** | ***SRSF2*** | ***NPM1*** | ***PTPN11*** |
|---|---|---|---|---|---|---|---|---|---|---|---|---|---|---|---|---|---|
| Taube et al.[21], Blood, 2022 | **131** | 35% | 24% | 18% | 14% | 13% | 2% | 10% | 4% | 4% | 5% | 5% | 3% | 2% | 2% | 3% | 0% |
| Zhang et al.[22], Cancer Gene Ther, 2019 | **76** | 32% | 5% | 30% | 21% | 16% | 13% | 0% | 1% | 0% | 11% | 7% | 1% | 3% | 0% | 1% | 1% |
| Su et al.[23], Oncotarget, 2018 | **81** | 16% | 12% | 18% | 15% | 10%* | 20% | 5% | 8% | 2% | 5% | 9% | N.D. | N.D. | 4% | 2% | N.D. |
| Konstandin et al.[24], Blood Adv, 2018 | **48** | 35% | 42% | 30% | 15% | 23% | 13% | 15% | 0% | 6% | 2% | 4% | 2% | 2% | 8% | 0% | 0% |
| Ahn et al.[25], Ann Hematol, 2016 | **51** | 14% | 8% | 14% | 8% | 8%* | N.D. | 6% | 14% | 4% | N.D. | N.D. | N.D. | N.D. | N.D. | 4% | N.D. |
| Papaemmanuil et al.[26], N Engl J Med, 2016 | **66** | 21% | 9% | 21% | 27% | 19% | N.D. | 8% | 6% | 0% | 6% | 2% | 0% | 5% | 0% | 3% | 5% |
| Fasan et al. [27], Leukemia, 2014 | **104** | 21% | 35% | 14% | N.D. | 3% | N.D. | N.D. | 7% | 14% | N.D. | N.D. | 6% | N.D. | N.D. | 2% | N.D. |
| **Overall (%)** (mutated/all) | **100%** **557/557** | **26%** 142/555 | **20%** 107/542 | **19%** 103/556 | **17%** 75/453 | **12%** 67/557 | **10%** 34/336 | **7%** 32/453 | **5%** 30/557 | **5%** 26/557 | **5%** 23/402 | **5%** 21/402 | **3%** 12/425 | **3%** 9/321 | **2%** 10/402 | **2%** 13/557 | **1%** 4/321 |

N.D. not determined; * *FLT3*$^{TKD}$ only, *FLT3*$^{ITD}$ N.D.

**Supplemental table 2A: Overlap of *TET2*$^{MUT}$ with *GATA2*$^{MUT}$ in *CEBPA*$^{DM}$ AML cases**

| Cohort | *CEBPA*$^{DM}$ cases (n) | *TET2*$^{WT}$ *GATA2*$^{WT}$ (n) | *TET2*$^{WT}$ *GATA2*$^{MUT}$ (n) | *TET2*$^{MUT}$ *GATA2*$^{WT}$ (n) | *TET2*$^{MUT}$ *GATA2*$^{MUT}$ (n) |
|---|---|---|---|---|---|
| Taube et al.[21], Blood, 2022 | 131 | 61 | 38 | 24 | 8 |
| Zhang et al.[22], Cancer Gene Ther, 2019 | 76 | 49 | 23 | 3 | 1 |
| Konstandin et al.[24], Blood Adv, 2018 | 48 | 15 | 13 | 16 | 4 |
| Ahn et al.[25], Ann Hematol, 2016 | 50 | 39 | 7 | 4 | 0 |
| Papaemmanuil et al.[26], N Engl J Med, 2016 | 66 | 48 | 13 | 4 | 1 |
| Fasan et al.[27], Leukemia, 2014 | 89 | 44 | 14 | 27 | 4 |
| **Total (n)** | **460** | **256** | **108** | **78** | **18** |
| ***TET2*$^{WT}$ cases (n) (%)** | **364** | **256 (70.3%)** | **108 (29.7%)** | – | – |
| ***TET2*$^{MUT}$ cases (n) (%)** | **96** | – | – | **78 (81.3%)*** | **18 (18.8%)*** |

* P=0.0103 (one-tailed Wilson/Brown Binominal test)

**Supplemental table 2B: Overlap of *TET2*$^{MUT}$ with *GATA2*$^{MUT}$ in all AML cases**

| Cohort | All cases (n) | *TET2*$^{WT}$ *GATA2*$^{WT}$ (n) | *TET2*$^{WT}$ *GATA2*$^{MUT}$ (n) | *TET2*$^{MUT}$ *GATA2*$^{WT}$ (n) | *TET2*$^{MUT}$ *GATA2*$^{MUT}$ (n) |
|---|---|---|---|---|---|
| Ahn et al.[25], Ann Hematol, 2016 | 388 | 326 | 12 | 49 | 1 |
| Papaemmanuil et al.[26], N Engl J Med, 2016 | 1540 | 1353 | 152 | 33 | 2 |
| **Total (n)** | **1928** | **1679** | **164** | **82** | **3** |
| ***TET2*$^{WT}$ cases (n) (%)** | **1843** | **1679 (91.1%)** | **164 (8.9%)** | – | – |
| ***TET2*$^{MUT}$ cases (n) (%)** | **85** | – | – | **82 (96.5%)*** | **3 (3.5%)*** |

* P=0.0491 (one-tailed Wilson/Brown Binominal test)

**Supplemental table 2C: Overlap of *TET2*$^{MUT}$ with *WT1*$^{MUT}$ in *CEBPA*$^{DM}$ AML cases**

| Cohort | *CEBPA*$^{DM}$ total cases (n) | *TET2*$^{WT}$ *WT1*$^{WT}$ (n) | *TET2*$^{WT}$ *WT1*$^{MUT}$ (n) | *TET2*$^{MUT}$ *WT1*$^{WT}$ (n) | *TET2*$^{MUT}$ *WT1*$^{MUT}$ (n) |
|---|---|---|---|---|---|
| Taube et al.[21], Blood, 2022 | 131 | 78 | 21 | *29* | 3 |
| Zhang et al.[22], Cancer Gene Ther, 2019 | 76 | 51 | 21 | 2 | 2 |
| Konstandin et al.[24], Blood Adv, 2018 | 48 | 25 | 3 | 16 | 4 |
| Ahn et al.[25], Ann Hematol, 2016 | 50 | 39 | 7 | 4 | 0 |
| Papaemmanuil et al.[26], N Engl J Med, 2016 | 66 | 40 | 21 | 5 | 0 |
| Fasan et al.[27], Leukemia, 2014 | 89 | 48 | 10 | 28 | 3 |
| **Total (n)** | **460** | **281** | **83** | **84** | **12** |
| ***TET2*$^{WT}$ cases (%)** | **364** | **281 (77.2%)** | **83 (22.8%)** | – | – |
| ***TET2*$^{MUT}$ cases (%)** | **96** | – | – | **84 (87.5%)**** | **12 (12.5%)**** |

** P=0.0081 (one-tailed Wilson/Brown Binominal test)

**Supplemental table 2D: Overlap of *TET2*$^{MUT}$ with *CSF3R*$^{MUT}$ in *CEBPA*$^{DM}$ AML cases**

| Cohort | *CEBPA*$^{DM}$ total cases (n) | *TET2*$^{WT}$ *CSF3R*$^{WT}$ (n) | *TET2*$^{WT}$ *CSF3R*$^{MUT}$ (n) | *TET2*$^{MUT}$ *CSF3R*$^{WT}$ (n) | *TET2*$^{MUT}$ *CSF3R*$^{MUT}$ (n) |
|---|---|---|---|---|---|
| Taube et al.[21], Blood, 2022 | 131 | 96 | 3 | 32 | 0 |
| Zhang et al.[22], Cancer Gene Ther, 2019 | 76 | 62 | 10 | 4 | 0 |
| Konstandin et al.[24], Blood Adv, 2018 | 48 | 26 | 2 | 17 | 3 |
| Ahn et al.[25], Ann Hematol, 2016 | - | N.D. | N.D. | N.D. | N.D. |
| Papaemmanuil et al.[26], N Engl J Med, 2016 | - | N.D. | N.D. | N.D. | N.D. |
| Fasan et al.[27], Leukemia, 2014 | - | N.D. | N.D. | N.D. | N.D. |
| **Total (n)** | **255** | **184** | **15** | **53** | **3** |
| ***TET2*$^{WT}$ cases (%)** | **199** | **184 (92.5%)** | **15 (7.5%)** | – | – |
| ***TET2*$^{MUT}$ cases (%)** | **56** | – | – | **53 (94.6%)** $^{n.s.}$ | **3 (5.4%)**$^{n.s.}$ |

N.D. not determined; $^{n.s.}$ P=0.3867 (one-tailed Wilson/Brown Binominal test)

**Supplemental table 2E: Overlap of *TET2*$^{MUT}$ with *ASXL1*$^{MUT}$ in *CEBPA*$^{DM}$ AML cases**

| **Cohort** | ***CEBPA*$^{DM}$ total cases (n)** | ***TET2*$^{WT}$ *ASXL1*$^{WT}$ (n)** | ***TET2*$^{WT}$ *ASXL1*$^{MUT}$ (n)** | ***TET2*$^{MUT}$ *ASXL1*$^{WT}$ (n)** | ***TET2*$^{MUT}$ *ASXL1*$^{MUT}$ (n)** |
|---|---|---|---|---|---|
| Taube et al.[21], Blood, 2022 | 131 | 98 | 1 | 28 | 4 |
| Zhang et al.[22], Cancer Gene Ther, 2019 | 76 | 72 | 0 | 4 | 0 |
| Konstandin et al.[24], Blood Adv, 2018 | 48 | 28 | 0 | 17 | 3 |
| Ahn et al.[25], Ann Hematol, 2016 | 50 | 42 | 4 | 4 | 0 |
| Papaemmanuil et al.[26], N Engl J Med, 2016 | 66 | 61 | 0 | 5 | 0 |
| Fasan et al.[27], Leukemia, 2014 | 89 | 47 | 11 | 26 | 5 |
| **Total (n)** | **460** | **348** | **16** | **84** | **12** |
| ***TET2*$^{WT}$ cases (%)** | **364** | **348 (95.6%)** | **16 (4.4%)** | – | – |
| ***TET2*$^{MUT}$ cases (%)** | **96** | – | – | **84 (87.5%)**** | **12 (12.5%)**** |

** P=0.0011 (one-tailed Wilson/Brown Binominal test)

**Supplemental table 3: CEBPA-mutated patients included in gene expression analysis (Beat AML study[19])**

| **Patient ID** | ***CEBPA*$^{MUT}$ (VAF)** | ***TET2*$^{MUT}$ (VAF)** | ***GATA2*$^{MUT}$ (VAF)** |
|---|---|---|---|
| AML_OHSU_2018_757 | Y108* (0.28) | - | - |
| AML_OHSU_2018_1049 | D80Sfs*78 (0.28) | - | Y322N (0.33) |
| AML_OHSU_2018_1159 | A47Cfs*61 (0.30) V314_L315dup (0.19) | - | - |
| AML_OHSU_2018_1370 | F82Sfs*78 (0.26) G340Cfs*84 (0.20) | M695Nfs*17 (0.30) Q1327* (0.26) | |
| AML_OHSU_2018_1396 | Q83Hfs*27 (0.20) | - | - |
| AML_OHSU_2018_1431 | K304_Q305insL (0.28) F82Lfs*28 (0.18) | - | N317H (0.22) |
| AML_OHSU_2018_1754 | L81Rfs*72 (0.40) | M823* (0.33) | - |
| AML_OHSU_2018_1848 | A30Pfs*130 (0.30) | - | - |
| AML_OHSU_2018_2073 | F77Sfs*83 (0.22) | - | - |
| AML_OHSU_2018_2113 | Y108* (0.19) | A1158V (0.19) | - |
| AML_OHSU_2018_2116 | T98Hfs*10 (0.32) | - | A318V (0.34) |
| AML_OHSU_2018_2311 | Q330* (0.26) | X1395_splice (0.35) C1289Tfs*75 (0.31) | - |
| AML_OHSU_2018_2322 | K313dup (0.27) S28Gfs*81 (0.24) | - | G320D (0.11) |
| AML_OHSU_2018_2630 | T98Hfs*10 (0.35) R297P (0.27) | Q758* (0.30) | - |
| AML_OHSU_2018_2713 | R35Gfs*125 | - | - |
| AML_OHSU_2018_2737 | K90Efs*18 (0.34) | - | - |

### Supplemental table 4: sgRNA sequences

| sgRNA | Sequence |
|---|---|
| sgTet2 (crRNA) | AAGGAGGCTGAAGAACAAGA |
| sgGata2 (crRNA) | TCTCTGGCGACGAGATGGCA |
| sgG2DHE_A | TGACGTAGCAAGCTGAGCGC |
| sgG2DHE_1 | TGGTCAGGTGGCGCTTATCA |
| sgG2DHE_2 | ATGGGCCTTCCGGAGCCCGT |
| sgG2DHE_3 | AGCCATCAGGCCCTGCAGCA |
| sgG2DHE_4 | GCAGGGGCTGTGAAGGCCCA |
| sgG2DHE_5 | GTATCCTGATGTGGTAAAGA |
| sgG2DHE_6 | TGCATGAATTCCGGTCTCAA |
| sgG2DHE_7 | TGTGAAACACCCCGAGCTTG |
| sgG2DHE_8 | GCTGTGCGGTGGGCAGAACG |
| sgG2DHE_9 | CACCCCCCAGGCAGTGGACA |
| sgG2DHE_10 | ATCATCTGCCAGCAGAGGCC |
| sgG2DHE_11 | CGCACAGCCTCCCTTAATTA |
| sgG2DHE_B | GAGCGACCTTTGCAGCAGCA |

### Supplemental table 5: PCR and qPCR primers

| **Target** | **Primer 1 (5' to 3')** | **Primer 2 (5' to 3')** | **Primer 3 (5' to 3')** |
|---|---|---|---|
| Cebpa wt-fl | GACTCCATGGGGGAGTTAGAG | GCCTTGGAAAGTCACAGGAG | - |
| Cebpa fl-ko | CCGCGGCTCCACCTCGTAGAAGTCG | CCACTCACCGCCTTGGAAAGTCACA | GTCCTGCAGCCAGGCAGTGTC |
| Cebpa p30 ki | CCGACTTCTACGAGGTGGAG | CTGTCGGCTGTGCTGGAAGA | - |
| Tet2 wt-fl-ko | GGCAGAGGCATGTTGAATGA | TAGACAAGCCCTGCAAGCAA | GTGTCCCACGGTTACACACG |
| Mx1-Cre | GCCTGCATTACCGGTCGATGCAACGA | GTGGCAGATGGCGCGGCAACACCATT | - |
| R26-Cre-ER | AAAGTCGCTCTGAGTTGTTAT | GGAGCGGGAGAAATGGATATG | CCTGATCCTGGCAATTTCG |
| R26-rtTA | AAAGTCGCTCTGAGTTGTTAT | GGAGCGGGAGAAATGGATATG | GCGAAGAGTTTGTCCTCAACC |
| MLL-AF9 ki | CTAGATCTCGAAGGATCTGGAG | ATACTTTCTCGGCAGGAGCA | - |
| miR30 | CAGAAGGCTCGAGAAGGTATATTGCTGTTGACAGTGAGCG | CTAAAGTAGCCCCTTGAATTCCGAGGCAGTAGGCA | - |
| Gata2 mRNA (for Cebpa$^{p30/-}$) | GCAGAGAAGCAAGGCTCGC | CAGTTGACACACTCCCGGC | - |
| Gata2 mRNA (for Cebpa$^{p30/p30}$) | ACAGGCCACTGACCATGAAG | AAGGGCGGTGACTTCTCTTG | |
| Gata2 V1 mRNA | CCGCTGCGAGTGGCC | GCCCGGATGGTGCGA | - |
| Gata2 V2 mRNA | GCCGCAGTCGGGCC | CTGCTCAGGCGCCACCT | - |
| Actb mRNA | AAGGAGATTACTGCTCTGGCTCCTA | ACTCATCGTACTCCTGCTTGCTGAT | - |
| Gapdh mRNA | AGAAGGTGGTGAAGCAGGCAT | CGGCATCGAAGGTGGAAGAGT | - |
| Gata2 V2 Bisulfite conv. DNA | GGAATTTTTTTAGTGGGATTTTAATAAG | ATACAATTTACTTACAATTTATCAACCC | - |
| Gata2 (TIDE genotyping) | TTTCCGGGTAACTTGCTGCT | TCAGGTGGTGAAGTGTCTGC | - |
| Tet2 (TIDE genotyping) | TGACTTTTCAGGGCTCGGTG | CGAGATCCCTCAACCATCGC | - |
| G2DHE (for ChIP-qPCR) | AATTCTGGTCAACCGCAAGC | TCTCGCATCCGTTACTTGCC | |

### Supplemental table 6: ShRNA sequences

| ShRNA | Identifier Mission® shRNA | Sequence of 97-mer (5' to 3') |
|---|---|---|
| ShGata2 A | TRCN0000085421 | TGCTGTTGACAGTGAGCGACCTGCAACACACCACCCGATATAGTGAAGCCACAGATGTATATCGGGTGGTGTGTTGCAGGGTGCCTACTGCCTCGGA |
| ShGata2 B | TRCN0000085419 | TGCTGTTGACAGTGAGCGACTCTACTACAAGCTGCACAATTAGTGAAGCCACAGATGTAATTGTGCAGCTTGTAGTAGAGGTGCCTACTGCCTCGGA |
| ShGata2 C | TRCN0000085418 | TGCTGTTGACAGTGAGCGACCCTGTAAATACAACCTTCTTTAGTGAAGCCACAGATGTAAAGAAGGTTGTATTTACAGGGGTGCCTACTGCCTCGGA |
| ShGata2 D | TRCN0000321390 | TGCTGTTGACAGTGAGCGACCGCCATTACTGTGAATATTTTAGTGAAGCCACAGATGTAAAATATTCACAGTAATGGCGGGTGCCTACTGCCTCGGA |

### Supplemental table 7: Antibodies for flow cytometry and FACS

| Antigen | Name | Manufacturer | Catalogue # | Fluorophore | Clone | Dilution |
|---|---|---|---|---|---|---|
| CD45.2 | PE Mouse Anti-Mouse CD45.2 | BD Pharmingen™ (BD Bioscience) | 560695 | PE | 104 | 1:200 |
| CD3e | CD3e Monoclonal Antibody | eBioscience™ (Thermo Fisher Scientific) | 15-0031-82 | PE-Cy5 | 145-2C11 | 1:400 |
| CD45R/B220 | CD45R (B220) Monoclonal Antibody | eBioscience™ (Thermo Fisher Scientific) | 15-0452-83 | PE-Cy5 | RA3-6B2 | 1:400 |
| Ly76/Ter119 | TER-119 Monoclonal Antibody | eBioscience™ (Thermo Fisher Scientific) | 15-5921-81 | PE-Cy5 | TER-119 | 1:400 |
| Ly76/Ter119 | TER-119 Monoclonal Antibody | eBioscience™ (Thermo Fisher Scientific) | 25-5921-82 | PE-Cy7 | TER-119 | 1:400 |
| Ly6G+Ly6C/Gr1 | APC Rat Anti-Mouse Ly-6G and Ly-6C | BD Pharmingen™ (BD Bioscience) | 553129 | APC | RB6-8C5 | 1:400 |
| Ly6G+Ly6C/Gr1 | Ly-6G/Ly-6C Monoclonal Antibody | eBioscience™ (Thermo Fisher Scientific) | **15-5931-82** | PE-Cy5 | RB6-8C5 | 1:400 |
| CD11b/Mac1 | FITC Rat Anti-CD11b | BD Pharmingen™ (BD Bioscience) | 553310 | FITC | M1/70 | 1:800 |
| CD11b/Mac1 | PE/Cyanine5 anti-mouse/human CD11b Antibody | Biolegend™ (NordicBiosite) | 101210 | PE-Cy5 | M1/70 | 1:800 |
| CD117/c-Kit | CD117 (c-Kit) Monoclonal Antibody | eBioscience™ (Thermo Fisher Scientific) | 47-1171-82 | APC eF780 | 2B8 | 1:200 |
| CD41a | CD41a Monoclonal Antibody | eBioscience™ (Thermo Fisher Scientific) | **11-0411-82** | FITC | eBioMWReg30 | 1:200 |
| Ly6A+Ly6E/Sca-1 | Ly-6A/E (Sca-1) Monoclonal Antibody | eBioscience™ (Thermo Fisher Scientific) | **45-5981-82** | PerCp-Cy5.5 | D7 | 1:200 |
| CD16+CD32/FcgRII/III | CD16/CD32 Monoclonal Antibody | eBioscience™ (Thermo Fisher Scientific) | **56-0161-82** | Alexa Fluor 700 | 93 | 1:100 |
